# Decoding primitive endoderm - epiblast interactions using mouse ICM embryoids

**DOI:** 10.1101/2024.12.28.630545

**Authors:** Beatrice F. Tan, Olivier J.M. Schäffers, Sarra Merzouk, Eric M. Bindels, Danny Huylebroeck, Joost Gribnau, Cathérine Dupont

**Author notes:** last author (Cathérine Dupont).

## Abstract

Stem cell-based embryo models are promising alternatives for investigating early embryogenesis. We introduce two distinct models to replicate the dynamics between extra-embryonic endoderm and epiblast during mouse embryonic development. Inducible *Gata6* (i*Gata6*) embryoid bodies (EBs), exclusively derived from i*Gata6* embryonic stem (ES) cells, are valuable for modeling the position-dependent development of the primitive endoderm. Inner cell mass (ICM) embryoids, conversely, efficiently formed by aggregating ‘wild-type’ and i*Gata6* ES cells, accurately and at a comparable pace simulate *in vivo* development from E3.5 to E7.5. Notably, ICM embryoids model cell sorting and through a rosette-like stage, the transition of the epiblast from naïve to primed pluripotency. Furthermore, the absence of extra-embryonic ectoderm-like cells in this model, directs both the epiblast and visceral endoderm towards an anterior developmental fate. As such, i*Gata6* EBs and ICM embryoids are powerful tools for advancing our understanding of cell fate decisions during early embryonic development in mice.

## Introduction

Preimplantation development in mice is marked by two cell fate decisions, each resulting in lineage segregations [1]. In the blastocyst, the first segregation takes place at embryonic day 3-3.5 (E3-E3.5) with the formation of the trophectoderm (TE) and inner cell mass (ICM). The second segregation then ensues in the ICM with the formation of the primitive endoderm (PrE, hypoblast) and epiblast. Mechanisms that operate during this second decision involve positional effects, cell sorting, and apoptosis. As development progresses, the PrE not only forms the parietal endoderm but also generates the visceral endoderm (VE), which surrounds the epiblast when the latter transitions from a naïve to primed pluripotent state. Intercellular communication between the PrE/VE and the epiblast, and the reciprocal interpretation of it, modulates the development of each of these two lineages. However, understanding the actors and underlying gene regulatory network changes during these phases of embryogenesis is hampered by complex, overlapping and redundant molecular mechanisms, along the poor accessibility of the small mouse embryo in the uterus.

Stem cell-based embryo models have become attractive alternatives to study early development of mammalian embryos, but they are not without limitations. Murine integrative embryo models such as the blastoid [2–4] and the ETX embryoids [5–10], which respectively model pre- and post-implantation development, fail to accurately replicate the *in vivo* developmental stages between E3-E5.5. Blastoids exhibit poor developmental potentials as the formation of their PrE (E3.5-E4) remains difficult and is dependent on various culture additives [2, 11]. ETX embryoids, conversely, are still in the assembly mode during that specific phase of development and are therefore most useful to model and study embryogenesis from E5.5 onwards. Furthermore, achieving high efficiency in both integrative embryo models poses significant challenges, impacting any further application. Several non-integrated stem cell-based embryo models have been described to partially mimic embryonic events between E3-E5.5. These models include embryoid bodies (EBs) generated from embryonic stem (ES) cells that are chemically or transgenetically manipulated to form PrE-like cells [11, 12], and mingling ES cells with Extra-Embryonic Endoderm (XEN) cells [5]. These models face limitations as well. Firstly, converting ES cells into XEN cells, when using a chemical process, is not ideal for understanding the communication between the epiblast and PrE. Secondly, converting all ES cells to PrE-like cells through *Gata4/6* overexpression does not accurately represent the cognate ICM. Thirdly, EBs formed from ES cells and XEN cells are also not ideal to model the ICM, as XEN cells are considered to be a differentiated cell type from the PrE [13].

Here, we detail the robust generation of ICM-resembling structures using two methods: aggregation of inducible *Gata6* Cherry-labeled (i*Gata6*) ES cells alone, termed mouse i*Gata6* EBs; and aggregation of ‘wild-type’ EGFP-labeled ES cells with i*Gata6* ES cells, termed mouse ICM embryoids. While i*Gata6* EBs aid in understanding the position-dependent development of the PrE, mouse ICM embryoids efficiently recapitulate epiblast and PrE dynamics from E3.5 onwards. These dynamics encompass cell sorting, pluripotency transitions, and the further development of the primed epiblast. Via immunostainings and scRNA-seq analysis, we compared the ICM embryoids with *in vivo* embryos, and validated decades of traditional research with these models. Furthermore, our scRNA-seq data revealed a dynamic and bi-directional interplay between the epiblast- and PrE-derived lineages affecting specific signaling pathways activity and accordingly developmental fate. ICM embryoids complement current embryo models by providing not only an efficient but also a potent tool for advancing our understanding of preimplantation development.

## Results

### An inducible CRISPRa system can robustly upregulate Gata6 mRNA levels in ES cells

To form a bi-potential ICM cell having the potential to form PrE and epiblast, we generated inducible (i*Gata6*) ES cell lines using a CRISPRa-system. For this, a CAG-driven rtTA cassette and a doxycycline-inducible dCas9VPR cassette were integrated in the TIGRE loci of mouse ES cells. Subsequently, a Cherry-labeled PiggyBac construct containing two guides that target the *Gata6* promoter was transfected into the ES cells (Fig. S1A).

After expansion of the constitutively Cherry-positive ES cell lines, the correctly targeted clones were tested for *Gata6* upregulation following docycyline induction (Fig. S1B, S1C). Expression levels of *Gata6, Sox17, Gata4, Pou5f1, Nanog* and *Sox2* in both i*Gata6*- and control *Actin*-EGFP ES cells were quantified following 6, 12, 24, 48 and 72h of doxycyline exposure. Compared to the control *Actin*-EGFP ES cells, the tested i*Gata6* ES cell line robustly upregulated *Gata6* transcription after 6h induction. At 24h, transcription of *Sox17* and *Gata4* became apparent indicating that the cells were further maturing into the PrE lineage. During this time frame, both the control and i*Gata6* ES cell lines showed downregulation of key genes associated with the pluripotency network, with the tested i*Gata6* ES cell line demonstrating more robust downregulation. (Fig. S1D).

### Position-dependent development of PrE

ICM cells of the early blastocyst (E3-E3.25) are uncommitted and have the potential to form either PrE or epiblast as they co-express *Nanog, Gata6* and *Pou5f1* [14, 15]. As development proceeds, ICM cells commit asynchronously to PrE or epiblast [16], resulting in a mixed population by E3.5 (Fig. 1A). Positional cues, together with signaling pathways, cell sorting and apoptosis, will guide lineage decisions within the ICM, leading to a mature PrE layer enveloping naïve epiblast cells at E4.5. The asynchronous developmental heterogeneity typically observed within the ICM complicates the understanding of underlying mechanisms.

**Figure 1.**
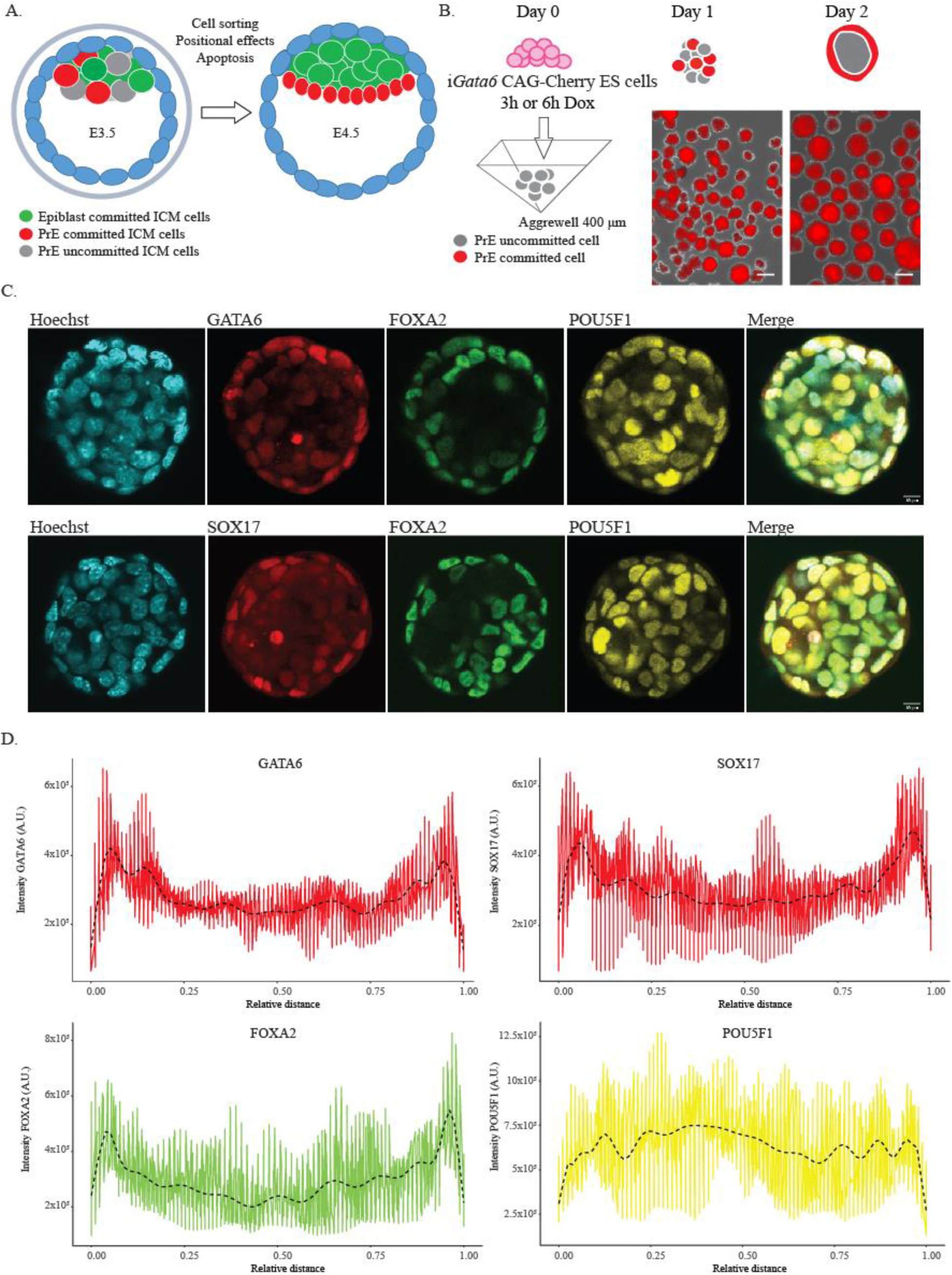
i*Gata6* embryoids model position-dependent development of the primitive endoderm. See also Figure S2 and S3A. A. The mouse E3.5 ICM is a heterogeneous cell mass that relies on various redundant cellular mechanisms to become organized at E4.5. B. Cherry-labeled i*Gata6* ES cells were briefly (3 to 6h) exposed to doxycycline (Dox) before being aggregated in Aggrewell plates (400 μm) in ETX medium (see Materials and Methods), scale bar 100µm. Primarily, cells located at the periphery of the iGata6 embryoids became fully committed to the PrE lineage. C. i*Gata6* EBs were collected on day 2 post-aggregation (following 3h Dox exposure) and assessed for PrE/VE and epiblast markers. In the top panel, i*Gata6* EBs stained for GATA6, FOXA2 and POU5F1. In the bottom panel i*Gata6* EBs stained for SOX17, FOXA2 and POU5F1, scale bar 10µm. D. Quantification of GATA6, SOX17, FOXA2 and POU5F1 intensity profiles measured across multiple line scans (*n*=2-4) per EB (*n*=10 per staining). Trend analysis is performed using orthogonal polynomial fitting (dashed line).

In mice, *Gata6* is expressed in all blastomeres of the 8-cell stage embryo [14, 17], but is not sustained in cells that do not commit to the PrE lineage [18]. To test the effect of positional cues in PrE development, i*Gata6* ES cells were briefly exposed for 3 to 6h to doxycycline to induce *Gata6* transcription, and subsequently aggregated to form i*Gata6* embryoid bodies (EBs) (Fig. 1B). The exposure time was varied to capture a bi-potent ICM cell state, with shorter exposure (3h) potentially better preserving the cell’s potential to form either epiblast or PrE. In contrast, a 6h exposure may already direct the ICM cell more decisively toward the PrE fate. GATA6 and other PrE transcription factors (SOX17 and FOXA2) were assessed after 48h to detect PrE-forming cells. After 48h, GATA6 and SOX17 were predominantly observed in the outer cells of i*Gata6* EBs, with reduced staining intensity detected in the inner cells, and – albeit less distinct –also for FOXA2 (Fig. 1C, Fig. S2A-B). Conversely, POU5F1 was more abundant in the inside of i*Gata6* EBs. Quantification of intensity profiles of each of these four transcription factors confirmed these observations (Fig. 1D). Longer exposure to doxycycline, also during the first 24h of aggregation (additional to the first 6h before aggregation), showed that the outer layer of the EBs was still more biased to form PrE compared to inner cells (Fig. S2C-D). These observations imply that the position of i*Gata6* cells within the EBs affects their commitment to the PrE-like lineage. Our findings support previous research, demonstrating the upregulation of the PrE marker PDGFRA in blastocoel-facing cells, while observing a corresponding downregulation in ‘inside’ ICM cells [14]. Overall, a brief induction of *Gata6* transcription led to the formation of uncommitted ICM cells displaying a position-influenced bias towards PrE formation.

*In vivo,* specification of the TE in the outer cells of the morula relies on the inhibition of the Hippo signaling pathway in response to the emergence of cell polarity. This inhibition prompts the upregulation of TE-specific genes [19, 20], while simultaneously suppressing the expression of *Sox2* [21, 22]. The Hippo pathway may similarly influence PrE maturation, as evidenced by the reduced contribution of *Tead4* null blastomeres to the PrE [23]. The i*Gata6* EBs offer a straightforward method to evaluate the impact of the Hippo pathway on PrE maturation. Therefore, we aggregated i*Gata6* ES cells (induced for 6h) and formed i*Gata6* EBs, but simultaneously exposed these to either a Hippo pathway activator (YAP-TEAD Inhibitor 1, 50nM-10µM) or a Hippo pathway inhibitor (LPA sodium salt, 500nM-5 µM). After 48h, we observed that markers indicative of PrE development (SOX17 and FOXA2), remained primarily at the EBs’ periphery in all conditions (Fig. S3A). This suggests that the position-dependent PrE development within i*Gata6* EBs does not rely on the Hippo pathway, or at least not solely. However, it’s essential to consider potential limitations such as suboptimal concentrations or activity of the small molecules used. Further molecular and genetic investigations are needed to clarify the role of the Hippo pathway and its interactions with other signaling pathways in regulating position-dependent development of the PrE.

### Cell allocation by sorting of iGata6 ES cells to the periphery of ICM embryoids

The observation that the ICM of the early blastocyst is composed of a random mixture of cells displaying varying levels of GATA6 or NANOG challenges the hypothesis that position would be the only factor that determines cell fate [17]. The ICM containing the aforementioned random mix of epiblast and PrE progenitors, are subsequently sorted such that cells with high *Gata6* and/or *Pdgfra* expression levels, line the blastocoel cavity and cover the *Nanog*-expressing ICM cells [14, 17]. Ultimately, cells wrongly positioned according to their developmental bias undergo apoptosis [14].

To test whether such operational cell sorting in the ICM could be modelled [14, 17, 24], aggregates were formed using a mixture of epiblast-biased *Actin*-EGFP and PrE-biased Cherry-labeled i*Gata6* ES cells (referred to as ICM embryoids). The labeling of the used ES cell lines was pivotal for characterizing their behavior throughout the development of the ICM embryoids. For each ICM embryoid, on average 20 ‘wild-type’ ES cells (*Actin*-EGFP) were mixed with 3 to 6 i*Gata6* ES cells that were briefly exposed (for 6h) to doxycycline before and, optionally, during the first 24h of aggregation (Fig. 2A). Most of such ICM embryoids showed i*Gata6* ES cells allocated to the periphery of ICM embryoids after 36h, although a small percentage of ICM embryoids also displayed incomplete or absent cell allocation configurations (Fig. 2B). To produce ICM embryoids presenting i*Gata6* ES cells almost exclusively at the periphery, the ratio of used ES cells needed to be carefully titrated. Insufficient numbers of i*Gata6* cells led to ICM embryoids failing to fully encapsulate *Actin*-GFP ES cells, while an excess resulted in the i*Gata6* cells also being found within the interior of the embryoids (Fig. 2B-C). The optimal ratio of cell numbers will vary depending on the proliferation rate of each ES cell line and should, therefore, be empirically determined. Following the optimization of cell numbers, ICM embryoids entirely enveloped by i*Gata6* cells could be obtained in up to 85% of the cases. As a result of positional cues and cell sorting, our ICM embryoids displayed GATA6 and SOX17 exclusively at the periphery, in contrast with i*Gata6* EBs produced exclusively from i*Gata6* ES cells (Fig. 2D; Fig. S3B-C).

**Figure 2.**
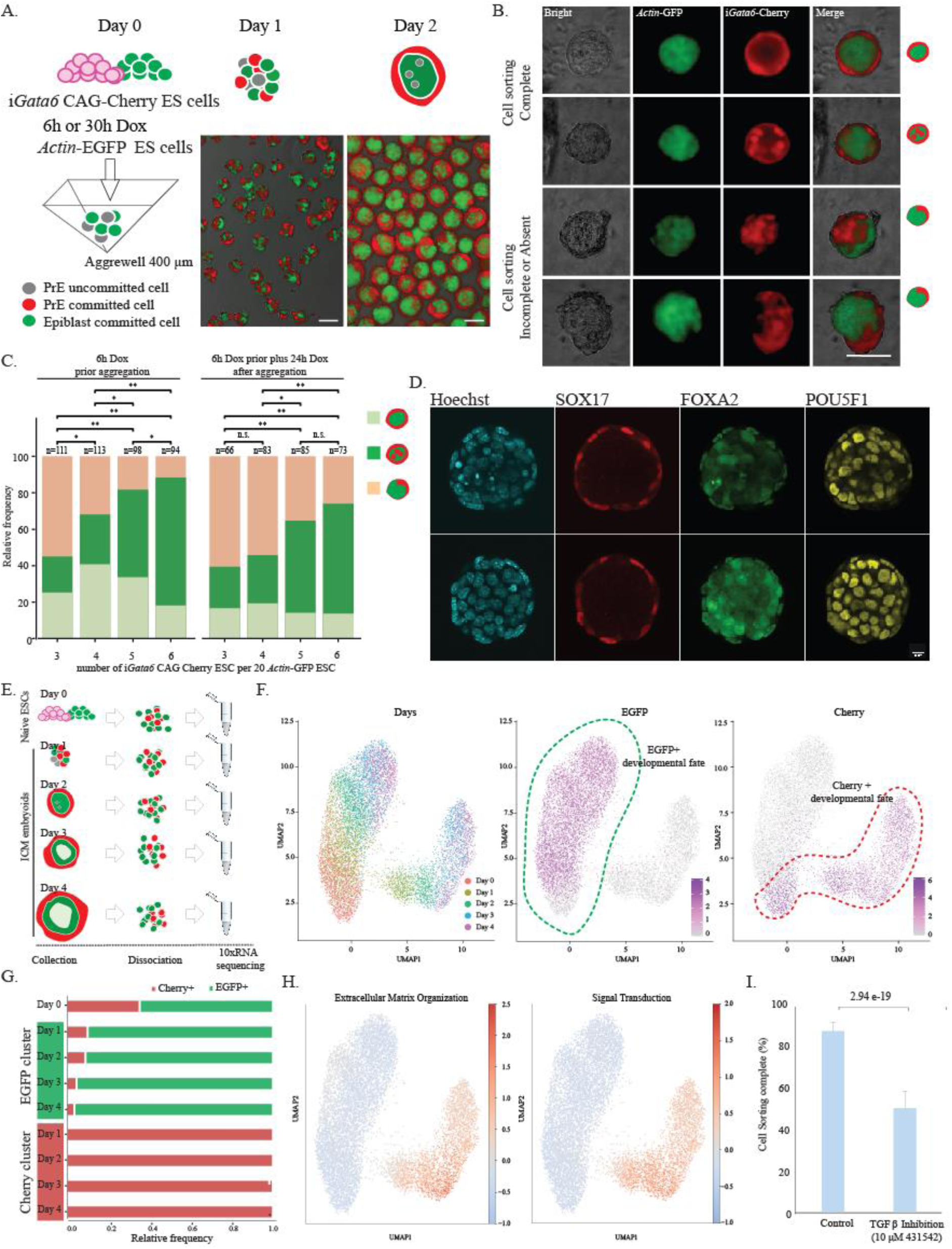
ICM embryoids model cell sorting of epiblast- and PrE-biased cells within the ICM. See also Figure S3B-C and Figure S4A-C. A. i*Gata6* ES cells were briefly exposed to doxycycline (Dox) before being aggregated with *Actin*-EGFP ES cells in Aggrewell plates (400 μm), and then collected at day 2, scale bar 100µm. B. Live-imaging of day 2 ICM embryoids showing complete, incomplete or absent cell allocation or sorting, brightfield overlapped with observed fluorescence (*Actin*-EGFP ES cells, CAG-Cherry ES cells (i*Gata6*)), scale bar 100 µm. C. Ratios of ICM embryoid configurations on day 2 post-aggregation (again following 6h of exposure to doxycycline). Examples of the different configurations are shown in B. Chi-square tests, adjusted for multiple testing with Bonferroni correction (* p < 0.05, ** p < 0.01), n = number of studied structures. D. Day 2 ICM embryoids stained for SOX17, FOXA2 and POU5F1 scale bar 10µm. E. Naïve ES cells (Day 0), and ICM embryoids at day 1, 2, 3 and 4 were collected, dissociated and processed for scRNA-seq. F. Integration of scRNA-seq data of naïve ES cells and ICM embryoids (day 1-4). UMAP plots showing composition of datasets and normalized expression of EGFP and Cherry. G. Evolution of percentage of Cherry-positive or EGFP-positive cells clustering within the EGFP or Cherry paths, respectively. H. UMAP plots showing enrichment of upregulated genes associated with the top Reactome terms between Cherry-positive cells of day 1-2 ICM embryoids compared to naïve ES cells. The top terms were selected based on their significant enrichment with the highest number of upregulated genes. I. ICM embryoids composed of wild-type (*Actin*-EGFP) and i*Gata6* ES cells display difficulties cell sorting following exposure to 10µM SB431542 (a TGFβ family signaling inhibitor). Bar plot depicting the average percentage of day 2 ICM embryoids displaying complete cell sorting with 95% confidence intervals, based on n=6 biological replicates per condition, statistical significance assessed using a chi-square test.

Single-cell RNA-sequencing (scRNA-seq) data from the ES cells in a naïve state, and following aggregation in ICM embryoids, showed that these two cell populations had two distinct developmental paths (Fig. 2E-F). A low percentage (approximately 8%) of Cherry-positive (i*Gata6*) cells initially adopted the epiblast fate and clustered with the EGFP (*Actin*-EGFP) cells. These cells, most likely represented i*Gata6* cells that did not fully mature towards the PrE-lineage as they were either positioned inside the ICM embryoids or did not express *Gata6* upon doxycycline induction. This percentage decreased as ICM embryoids further developed, suggesting that although induced i*Gata6* cells initially attempt to adopt the developmental fate as determined by their position, they exhibit a reduced competence to form the epiblast as development progresses (Fig. 2G). Conversely, no EGFP-positive (*Actin*-EGFP) cells clustered with the Cherry positive cluster. Functional enrichment analysis showed that many upregulated genes in derivatives of the i*Gata6* ES cells on day 2; are related to extracellular matrix (ECM) (e.g. Laminins, Collagens, Nidogens), cell adhesion and guidance proteins (e.g. Ephrins, Cadherins, Nectins) along signal transduction (Fig. 2H, Fig. S4A). These findings are in line with previous studies that implicate an important role for the ECM and cell interactions in cell sorting in the ICM [25, 26]. Additionally, signaling pathways are likely to contribute to this sorting mechanism as well. For instance, Disabled-2 (Dab2), identified as a crucial factor during ICM cell sorting [27] and detected in Cherry-positive cells from day 2 onwards (Fig. S4B), interacts with integrins and various signaling pathway receptors and adaptors [28–32]. Since TGFβ family ligands have been reported to increase the binding of DAB2 to integrins [33], we questioned whether cell sorting could be perturbed by blocking TGFβ family signaling. As expected, our day 2 ICM embryoids displayed a reduced cell sorting efficiency compared to control ICM embryoids following exposure to TGFβ inhibitor SB431542 (Fig. 2I, Fig. S4C). Given the importance of TGFβ family signaling in determining the cell fate of both the epiblast and the PrE, it may however not be excluded that the observed cell sorting perturbations are secondarily to suboptimal cell lineage specification. Nonetheless, these results show that ICM embryoids can be an essential tool to assess genes and signaling pathways that affect cell allocation by active cell sorting principles.

### Naïve epiblast cells gradually transition into a primed state of pluripotency within ICM embryoids

The E3.5 ICM displays a “salt and pepper” distribution of epiblast and PrE committed cells, which from E4.5 onwards become sorted with the PrE covering the naïve epiblast [17]. Day 1 ICM embryoids exhibited some cell sorting, which appeared to be complete by day 2. When ICM embryoids were allowed to develop beyond day 2, the formation of an epithelial-like epiblast surrounded by a VE-like cell layer became apparent (Fig.3A). Immunostainings of ICM embryoids at days-3 and -4 showed pluripotent (POU5F1) cell masses surrounded by an endoderm-like cell layer (SOX17) (Fig.3B). Differential expression analysis of scRNA-seq data from our ICM embryoids at different time points revealed that many lineage-specific genes were among the top differentially expressed genes (DEGs), including *Klf2* and *Zfp42* in naive ES cells, *Gata6* and *Dab2* in Cherry-positive cells, and *Pou3f1* and *Otx2* in EGFP-positive cells (Table S1). Enrichment analysis of these DEGs on scRNA-seq data from *in vivo* E3.5-E7.5 embryos [34] showed that both the epiblast- and PrE-like cells developed towards tissues resembling undifferentiated epiblast of E6.5-E7.5 and endoderm cell types, respectively (Fig. 3C-E, Fig. S4D-F).

**Figure 3.**
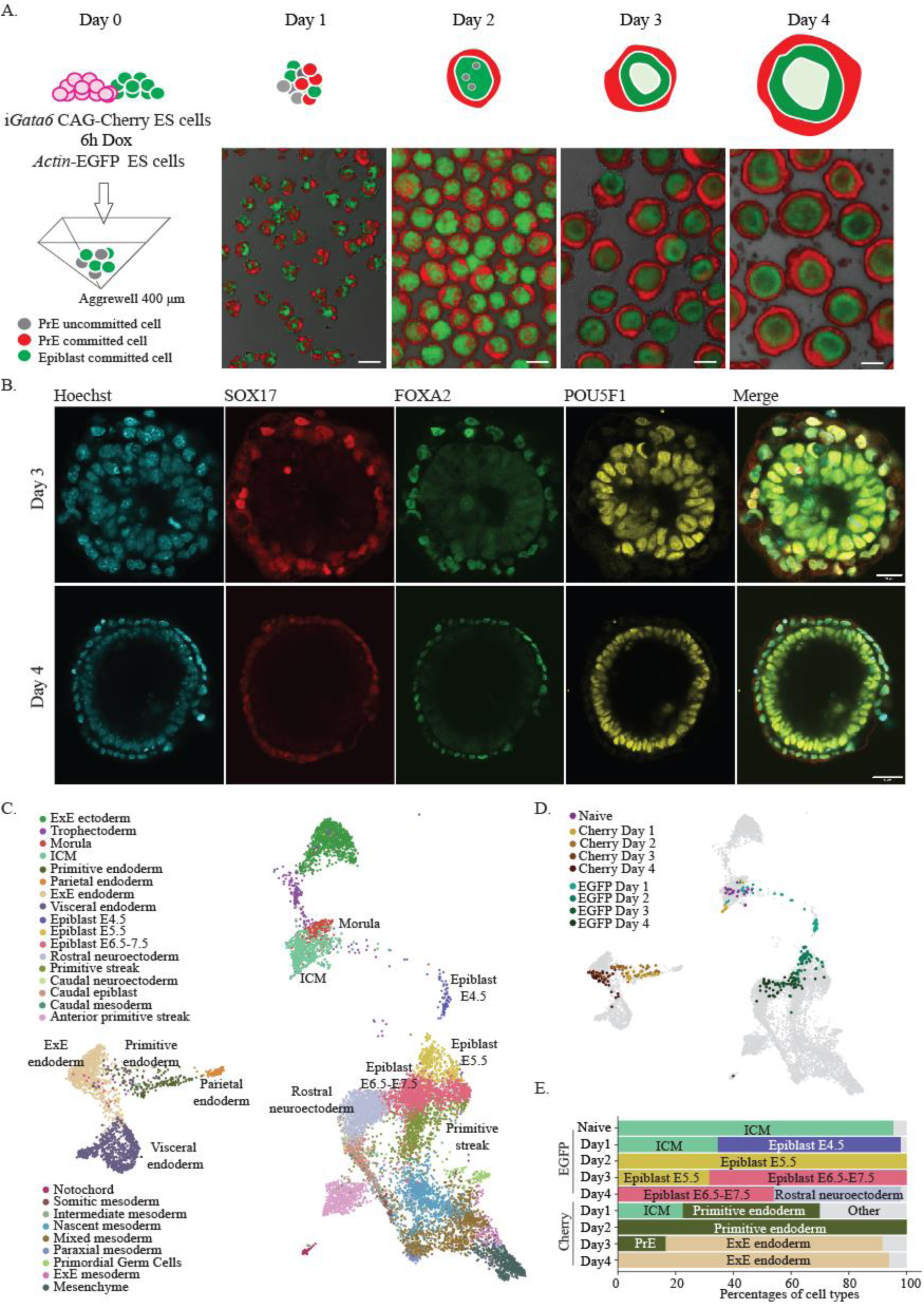
ICM embryoids display developmental progression resembling *in vivo* developmental stages. See also Figure S4D-F. A. Procedure for making ICM embryoids composed of wild-type (*Actin*-EGFP) and Cherry-labeled i*Gata6* ES cells. i*Gata6* ES cells (briefly induced during 6h), are sorted to the periphery of most ICM embryoids on day 2. Epiblast formation can be morphologically observed from day 3 onwards, scale bar 100µm. B. On day 3 and day 4, ICM embryoids stained for SOX17 and FOXA2 (both marking PrE-derived cells) and POU5F1 (marking the epiblast). Scale bars day 3 and day 4, 20µm and 40µm; respectively. C. UMAP showing clustering of *in vivo* cells from morula to E7.5 gastrulation in embryonic and extraembryonic lineages colored according to the cell type labels from [34]. Cell types relevant for the comparison to the ICM embryoids are labeled as well. ICM, inner cell mass. ExE, extraembryonic. D. UMAP highlighting the *in vivo* cells most similar to those from the ICM embryoids. For each group of ICM embryoid cells (Naive, EGFP-/Cherry-positive per day), the 50 embryo cells with the highest of enrichment of its respective markers were labeled. E. Barplot showing the percentage of each *in vivo* cell type annotation of the top cells per ICM embryoid cell cluster. Cell types with at least 10 occurrences are individually labeled (and colored according to the legend in C), while other cell types are grouped under ‘other’. PrE, primitive endoderm.

The developmental trajectory of EGFP-positive cells, as determined through pseudotime and RNA velocity analysis, corresponded with the sequential stages of epiblast development observed in ICM embryoids collected over consecutive days (Fig. 4A, Fig. S5A). The progression of the naïve epiblast towards its primed pluripotent state was, in both ICM embryoids and in natural embryos, assessed via a selection of transcriptional markers (Fig. 4B-D, S5B-C). Epiblast cells in day 1 ICM embryoids and E4.5 embryos, expressed solely naïve pluripotency transcription factors such as *Zfp42, Klf4, Esrrb* and *Tbx3* (Fig. 4B-D, Fig. S5B-C). During further development, both epiblast cells of day 2 ICM embryoids and E5.5 embryos, expressed mostly only markers of primed pluripotency (*Fgf5, Otx2, Pou3f1, Zic2* and *Zic5*). This implied that the intermediate stage of pluripotency, also denoted as naïve-primed intermediate [35], poised [36] or formative [37], occurred between day-1 and -2 in ICM embryoids (Fig. 4D). Therefore, day 2 ICM embryoids resembled the late rosette stage occurring *in vivo* at E5.25 [38]. Immunostaining for Phalloidin and POU5F1, revealed lumen emergence in day 2 ICM embryoids, further confirming the staging of these embryoids (Fig. 4E, Fig. S5D). Smaller day 2 ICM embryoids displayed a singular central lumen as observed in *in vivo* rosette stage embryos (E5.25), while multiple lumens could be observed in larger day 2 ICM embryoids (Fig. 4E, Fig. S5D). At day-3 and -4, the pluripotent stem cells had clearly matured into a primed state without expression of naïve pluripotency markers, matching transcriptomic data from E6.5-E7.5 *in vivo* epiblast (Fig. S5C). Moreover, the epiblast-like cells in day 4 ICM embryoids also expressed genes associated with rostral neuroectoderm, as confirmed by an enrichment of the rostral neuroectoderm signature at that stage (Fig. 3E, Fig. S5E). The epiblast of day 4 ICM embryoids, however, most likely adopted an anterior epiblast fate, a cell type that was not separately annotated in the embryo scRNA-seq dataset that was used for the analysis, which during further culture normally would have formed rostral neuroectoderm.

**Figure 4.**
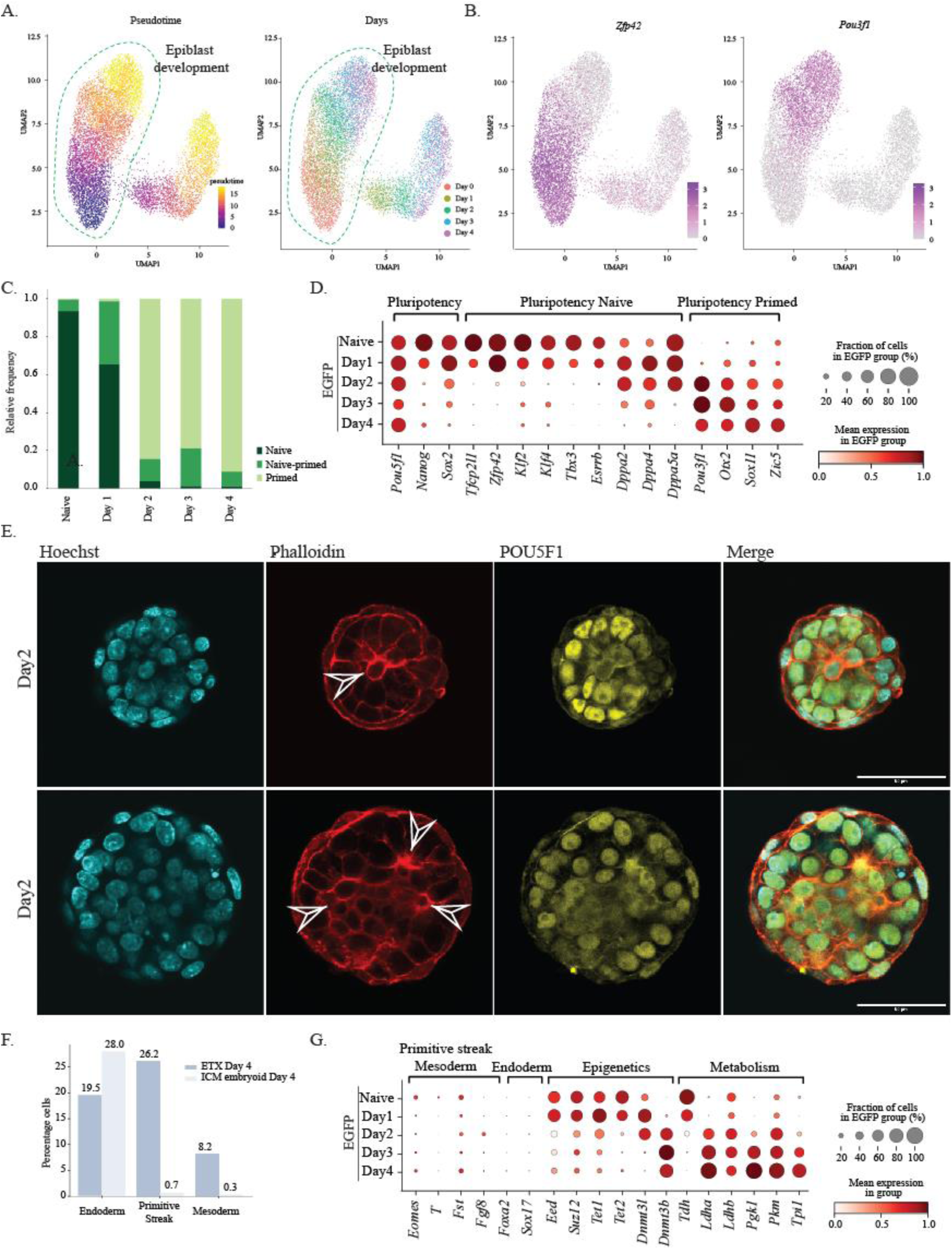
Naive to primed pluripotency transition occurs between day 1 and day 2 in epiblast-like cells of ICM embryoids. See Figure S5. A-B. Integration of scRNA-seq data of naïve ES cells and ICM embryoids (day 1-4) UMAP plots colored by relative pseudotime predicted by Monocle3, composition of datasets and normalized expression of *Zfp42* and *Pou3f1*. C. Bar chart displaying the percentage of EGFP-positive cells showing transcripts of naïve markers (*Zfp42* and/or *Klf4*) and/or primed markers (*Otx2* and/or *Pou3f1*). D. Dot plot showing expression of naïve and primed markers in EGFP-positive populations over time in ICM embryoids. Color indicates normalized gene expression, while dot size reflects the percentage of cells expressing the gene. E. Lumen formation in day 2 ICM embryoids stained for Phalloidin and POU5F1 (arrowheads indicate lumen formation), scale bar 50µm. F. Bar chart displaying the percentage of EGFP-positive cells with transcripts for endoderm (*Foxa2, Sox17*), primitive streak (*Fgf8* and *Fst*) and mesoderm (*T, Eomes*) formation in day 4 ICM embryoids and day 4 ETX embryoids. G. Dot plot showing normalized expression of markers for endoderm and mesoderm/primitive streak along markers for epigenetic profile and metabolism in ICM embryoids.

Failure to re-upregulate *Nanog* and the absence of primitive streak-associated factors such as *Eomes, Brachyury/T* and *Fst* in day 4 ICM embryoids further validated the anterior fate of the epiblast (Fig. 4F). This developmental fate is likely due to the absence of ExEc in ICM embryoids as opposed to natural embryos and ETX embryoids, which typically plays a role in promoting gastrulation in the posterior-proximal region of the cylindrical epiblast (Fig. 4F). The progression from naïve to primed pluripotency in our ICM embryoids, involved not only alterations in the expression of transcription factor encoding genes, but also encompassed shifts in energy production and modifications in the epigenetic landscape (Fig. 4G). These modifications adhered to scRNA-seq data from *in vivo* embryos (Fig. S5C). Naïve epiblast cells use both glycolysis and oxidative phosphorylation for energy production, whereas primed epiblast cells solely rely on glycolysis [39]. As such, downregulation of oxidative phosphorylation/mitochondria genes (*Tdh*) and upregulation of glycolytic genes (*Pgk1, Pkm, Tpi1, Ldha, Ldhb*) could be observed as ICM embryoids transitioned from naïve to primed pluripotency. Also, DNA in naïve pluripotent stem cells is hypomethylated compared to the primed state, with transcriptional repression relying more on histone modifications [40–42]. In *in vivo embryos*, this is associated with a specific expression profile of *Dnmt3a, Dnmt3b, Dnmt3L* and PRC2 complex subunits [43]. In ICM embryoids, *Dnmt3L* was, similar to *in vivo* embryos, highly expressed in the naïve epiblast while *Dnmt3a* and *Dnmt3b* were upregulated when primed pluripotency was established. Furthermore, and also similarly to *in vivo* embryos, fewer transcripts of DNA-demethylation enzymes (*Tet1* and *Tet2*) and PRC2 complex subunits (*Suz12* and *Eed*) could be observed after the epiblast had transitioned towards primed pluripotency.

### The PrE matures into distal and/or anterior visceral endoderm cell types in the absence of extra-embryonic ectoderm

While the developmental path of i*Gata6* cells, as predicted by pseudotime and RNA velocity analysis (Fig. 5A, Fig. S5A), aligned with the early days of ICM embryoid formation, i*Gata6* cells from day 3 and 4 ICM embryoids exhibited greater diversity in developmental stages. Louvain clustering was performed to obtain clusters representing the different stages of endoderm development (Fig 5A). In ICM embryoids, the PrE had matured by day 1, as evidenced by the expression of *Sox17* and *Gata4* (Fig. 5A-B). As development proceeds, the PrE that envelops the epiblast and the ExEc, will form the VE. By analyzing transcripts of *Spink1* and *Ttr* [44, 45], it became evident that the PrE-derived lineages observed in day 2 ICM embryoids underwent progressive development into VE rather than adopting a parietal endoderm fate (Fig. 5B). The presence of *Eomes* and *Fgf5* transcripts highlighted that the VE rather formed embryonic VE (emVE) than extra-embryonic endoderm (ExEn) [46, 47] (Fig. 5B). The comparison of scRNA-seq data of the ICM embryoids to *in vivo* embryos, however, showed that the PrE-derived cells in our ICM embryoids mostly resembled ExEn rather than emVE cell types (Fig. 3C-E). This discrepancy may be attributed to the limited characterization of PrE-derived lineages in the *in vivo* dataset used. To better profile the PrE-derived cells in our ICM embryoids, we used another *in vivo* dataset that focused on characterizing the PrE-derived lineages [48] (Fig. S6A-B). As expected, DEGs from Cherry-positive cells in day 1 to day 4 ICM embryoids were initially enriched in PrE cells of day 4.5 blastocysts and subsequently in emVE cells of day 6.5-7.5 *in vivo* embryos as the culture progressed (Fig. S6C).

**Figure 5.**
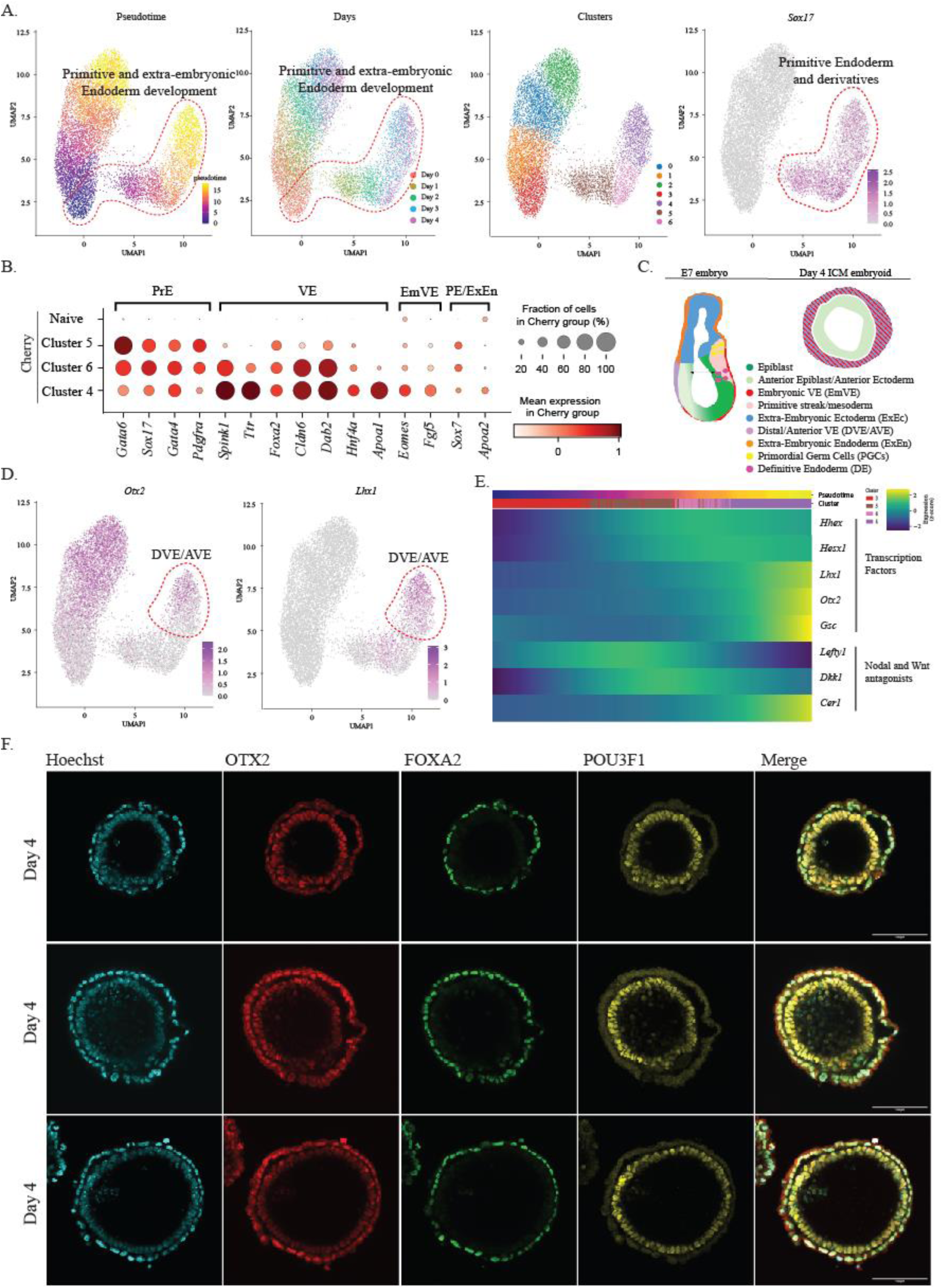
The default developmental state of the PrE is the DVE/AVE. A. UMAP plots colored by relative pseudotime showing primitive and extra-embryonic endoderm development in the Cherry-positive clusters, composition of datasets, clusters identified using Louvain and normalized expression of *Sox17.* B. Dot plot showing normalized expression of endoderm markers in naive ES cells and Cherry clusters in ICM embryoids. Color indicates normalized gene expression, while dot size reflects the percentage of cells expressing the gene. C. Schematic representation of day 4 ETX and ICM embryoids. D. UMAP plots displaying normalized expression of *Otx2* and *Lhx1*, DVE/AVE markers. E. Heatmap showing expression of several transcription factors and Nodal and Wnt antagonists in Cherry-positive cells ordered along pseudotime. The cells are annotated above the heatmap according to the pseudotime and clusters shown in A, z-score of normalized expression is plotted. F. Day 4 ICM embryoids stained for OTX2, FOXA2 and POU3F1, OTX2 in the VE marks DVE/AVE, scale bar 100µm.

The analysis of the pseudotime trajectory of PrE-derived cells in our ICM embryoids revealed the presence of several key DVE/AVE markers (*Otx2, Hhex, Lhx1, Dkk1, Cer1* and *Lefty1,* [49–59]) at the endpoint of the developmental trajectory (Fig. 5A-E). This suggests that the formation of the DVE/AVE is the default developmental state in ICM embryoids. Supporting this notion, immunostained day 4 ICM embryoids often revealed a distinct accumulation of OTX2 in the emVE-like cell layer (Fig. 5F). Since previous studies have suggested that the ExEc limits the number of DVE/AVE cells observed in *in vivo* embryos [60], the observations that the emVE in the ICM embryoids (lacking ExEc) had a DVE/AVE character seems quite logical (Fig. 5C).

### Reciprocal communication between epiblast- and PrE-derived lineages

The developmental trajectory of PrE- and epiblast-derived lineages within ICM embryoids was dictated by various intercellular interactions, as predicted by CellPhoneDB [61], including ligand-receptor, transmembrane protein and extracellular matrix (ECM) interactions (Fig. S7A-C). While both cell populations engaged in this communication, PrE-derived cells emerged as the predominant mediators of these interactions within ICM embryoids (Fig. 6A, Fig. S7A). Additionally, the majority of ECM-related transcripts, which constitute the basement membrane [62], were produced by PrE-derived lineages (Fig. 6B). Our analysis detected a variety of transmembrane proteins expressed in both epiblast- and PrE-derived lineages; including Ephrin-Ephrin receptors, Integrins, Cadherins, Nectins, Teneurins, Flirtuins, Neurexins, and Latrophilins (Fig. S7B-C, Fig. S8A). Besides those, also various ligand-receptor interactions could be observed. Given the diverse array of intercellular interactions, all of which have the potential to affect signaling pathway activity, it is crucial to refrain from assuming pathway activity solely based on the detection of ligand-receptor interactions. Therefore, we also examined a subset of effector genes whose transcriptional upregulation serves as a marker of pathway activity (Fig. S8B). Expression of these effector genes was averaged to summarize the pathway activity in the lineages during ICM embryoid development. Additionally, the expression of specific ligands and receptors was examined to further validate our findings.

**Figure 6.**
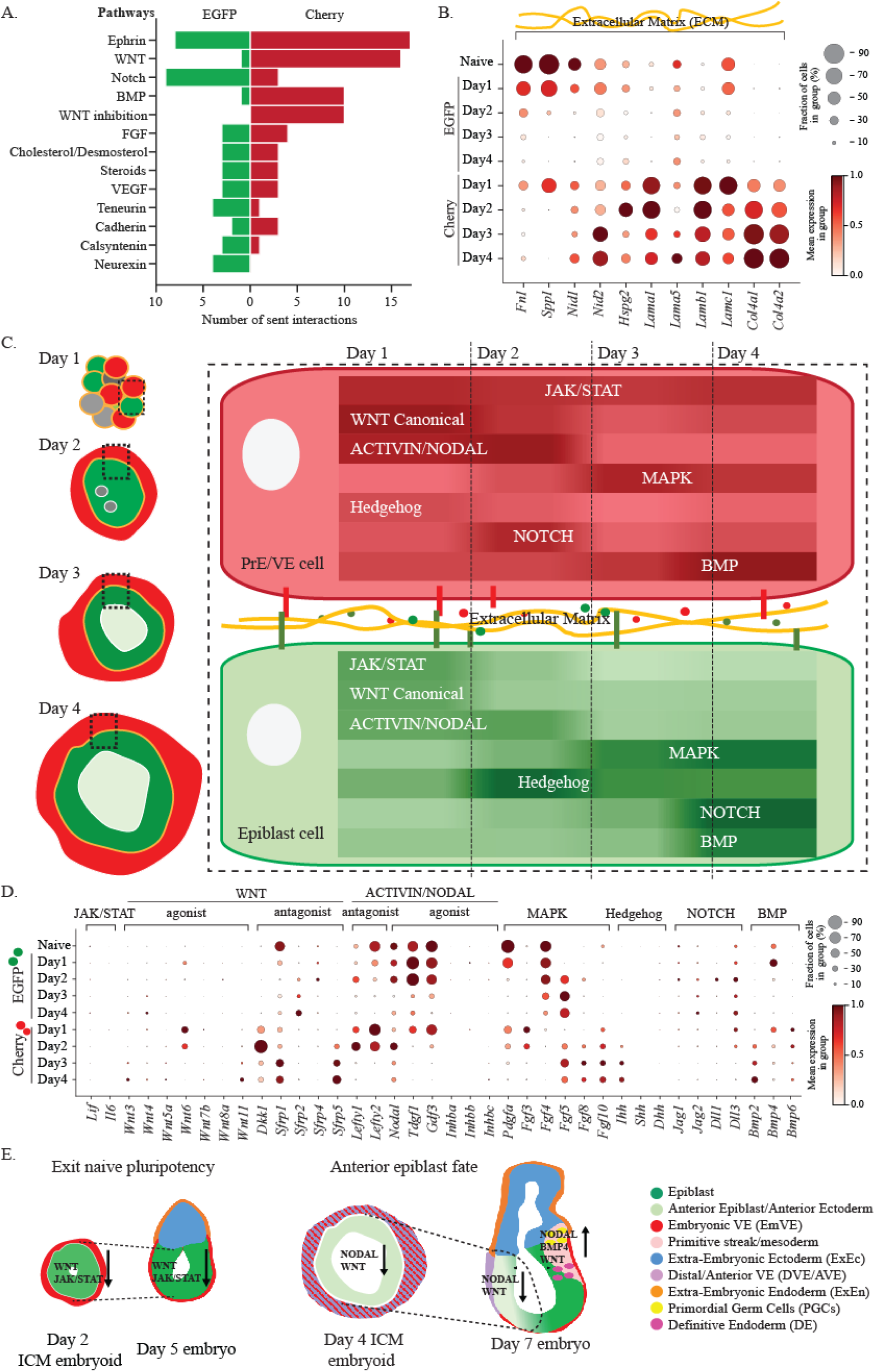
The anterior developmental fate is determined by the absence of the ExEc and the interactions between the PrE and the epiblast. See also Figure S7-S8. A. Barplot showing the number of active interactions per pathway originating from the EGFP (green) or Cherry lineage (red) as predicted by CellphoneDB. The bars represent the number of significant interactions in at least one day. Only pathways active in at least one of the lineages are shown. B. Dot plot showing normalized expression of extracellular matrix transcripts in naïve ES cells and in ICM embryoids. Color indicates normalized gene expression, while dot size reflects the percentage of cells expressing the gene. C. Schematic representation of pathway activity based on expression of assessed effector genes in Cherry positive and EGFP positive lineages in ICM embryoids D. Dot plot showing normalized expression of ligand transcripts implicated in various signaling pathways in naïve ES cells and ICM embryoids. E. Schematic representation displaying similar pathway activity in the epiblast of day 2 versus day 5 embryos and day 4 ICM embryoids versus the anterior epiblast of day 6.5-7 egg cylinder stage embryos.

According to our assessment of effector genes, the exit from naïve pluripotency in the epiblast of day 2 ICM embryoids was accompanied by a decrease of JAK/STAT (*Socs3*, *Stat3, Cd9*) [63, 64] and canonical WNT (*Axin2, Tbx3*) [65–67] signaling alongside an increase in MAPK/ERK signaling (via *Fos*, *Egr1*, *Jun*) [68] (Fig. 6C, Fig. S8B). These observations generally matched with the expression data from the corresponding *in vivo* epiblast (Fig. S8D) and showed an inverse relationship with the pathway activity observed in naïve ES cells [69–74]. The PrE-derived lineages in day 2 ICM embryoids also exhibited decreased canonical WNT- and increased MAPK/ERK-signaling pathway activity. Conversely to the epiblast-derived cells, the PrE-derived lineages maintained JAK/STAT3 signaling activity (Fig. 6C, Fig. S8B). Since the ICM embryoids expressed neither *Lif* nor *Il6* (Fig. 6D), another unspecified interaction must have maintained JAK/STAT3 pathway activity in the PrE-derived lineages. Repression of WNT signaling during the exit from naïve pluripotency in ICM embryoids between day 1 and day 2, could at least be partially attributed to the expression of WNT antagonist encoding gene *Dkk1* [75] expressed by the formed PrE-like cells (Fig. 6D). Both the epiblast- and PrE derived lineages in the ICM embryoids independently produced growth factors and receptors that fostered MAPK/ERK activity within their own, but also counterpart, lineage (Fig. 6D, Fig. S8C). Among these factors, and similarly as described in natural embryos, the epiblast’s transcription of *Fgf4* and *Pdgfa* in day 1 ICM embryoids supported PrE maturation [76–79] and its survival [80, 81], respectively (Fig. S8E-F). Other observed *Fgf* transcripts such as *Fgf3*, *Fgf5*, *Fgf8* and *Fgf10* also supported general MAPK activity in both lineages and derivatives (Fig. 6D). In summary, whereas naïve epiblast cells of day 1 ICM embryoids supported the formation of PrE-like cells, formed PrE-derived lineages subsequently instigated the epiblast’s exit from naïve pluripotency by repressing WNT-signaling.

Based on the expression of effector genes *Lefty1, Lefty2*, and *Nodal*; ACTIVIN/GDF3/NODAL signaling appeared to decrease from day 3 onwards (Fig. 6C, Fig. S8B). The absence of *Activin*-along the detection *Gdf3-* and *Nodal*-transcripts in the ICM embryoids (Fig. 6D), mirrored the gene expression patterns observed in late blastocyst-stage natural embryos (E4.5-E5) (Fig. S8E). According to expression levels of *Axin2* and *Tbx3*, also WNT signaling remained low in all lineages after day 2 (Fig. 6C, Fig. S8B). The further inhibition of WNT signaling beyond day 2 was possibly facilitated by SFRP1 and SFRP5 [75], abundantly expressed by the PrE-derived lineages beyond day 2 (Fig. S8A-B, Fig. 6D). As deduced from BMP (-SMAD) responsive genes *Id1, Id2* and *Id3* [82] expressed in our ICM embryoids, BMP signaling incremented as development proceeded (Fig. 6C, Fig. S8B). Our ICM embryoids primarily expressed *Bmp2*, *Bmp4* and to a lesser extent *Bmp6* (Fig. 6D). Whereas *Bmp4* was exclusively expressed by naïve ES cells and day 1 ICM embryoids, *Bmp2* and *Bmp6* were expressed by the VE at later stages.

Insufficient maintenance of NODAL or exposure to BMP4 in the primed epiblast of mouse embryos hinders the upregulation of WNT signaling necessary for primitive streak formation [83–89]. In natural embryos, the ExEc secretes BMP4 and aids in sustaining NODAL signaling in the epiblast, which together induce WNT pathway activity in the posterior epiblast necessary for the formation of the primitive streak (Fig. 6E) [90, 91]. The absence of NODAL, BMP4 and consequently WNT signaling pathway activity in the epiblast of our day 3 and day 4 ICM embryoids is at least partially attributed to the absence of the ExEc.

## Discussion

The formation of the epiblast and PrE within the ICM depends on positional cues and cell sorting dynamics. The epiblast subsequently transitions from a naïve to primed pluripotent state before acquiring an anterior-posterior axis during further development. However, studying this developmental stage is challenging due to the fragility and small size of the embryo during implantation, making its dissection from the uterus without damage difficult, and the limitations of current murine integrative embryo models like blastoids [2–4] and ETX embryoids [5–10] with regard to modeling embryonic development between E3 and E5.5 and their low assembly efficiency.

Here, we propose i*Gata6* EBs and ICM embryoids as additional robust models for studying this phase of development. Forming EBs from solely i*Gata6* ES cells, a position-dependent development of the PrE at the outer rim of i*Gata6* EBs can be studied. Although our study did not provide conclusive evidence regarding the impact of the Hippo signaling pathway on PrE development, further investigation is underway to address this question. ICM embryoids, conversely, formed from aggregated *Actin*-EGFP and Cherry labeled i*Gata6* ES cells, can be used to study and model cell sorting dynamics observed in the ICM of blastocyst-stage embryos. Epiblast-biased cells in ICM embryoids transitioned from naïve to primed pluripotency between day 1 and day 2 passing through a rosette-like stage (E5.25). During further development the epiblast-biased lineages correlated at day 3 and day 4 with epiblast adopting an anterior developmental fate. Our comparative analysis with *in vivo* scRNA-seq data unveiled that the PrE-biased lineages within ICM embryoids developed into an emVE-like cell type, characterized by features reminiscent of both DVE and AVE. The observation that emVE adopts a DVE/AVE character is in accordance with data from *in vivo* embryos, where removal of the ExEc soon after implantation leads to emVE displaying DVE/AVE characteristics [60]. Our analysis also highlighted reciprocal signaling (via growth factors, transmembrane interactions and extracellular matrix proteins) between the PrE- and the epiblast-derived lineages of the ICM embryoids. Specifically, the PrE-biased lineages in day 1 and day 2 ICM embryoids supported the exit from naïve pluripotency primarily by producing extracellular matrix proteins and antagonizing WNT-signaling. The epiblast, conversely, promoted the maturation and survival of the PrE during these stages through the expression of *Fgf4* and *Pdgfa*. After the initial decrease in WNT- and JAK/STAT-signaling, there is a subsequent decline in NODAL/GDF3 signaling over the following days. While BMP signaling seemed to increase, this was mostly attributable to BMP2/BMP6 and not BMP4. The observed pathway activity in day 3 and day 4 ICM embryoids can logically be attributed to the absence of the ExEc in the ICM embryoids. In *in vivo* embryos, the ExEc maintains NODAL signaling in the posterior epiblast and produces BMP4, both necessary for the upregulation of WNT signaling in the posterior epiblast (Fig. 6E). This process ultimately results in the formation of the primitive streak. The activity of these pathways in the anterior epiblast in *in vivo* embryos are inversely correlated with those in the posterior epiblast. Consequently, the epiblast of ICM embryoids logically resembles the anterior epiblast. Our findings thus contrast with other VE/epiblast models, which are capable of forming mesoderm [11]. The mesoderm formation observed in these models is most likely due to WNT signaling stimulating ligands in the culture medium necessary for the formation of these models.

The default developmental state of the PrE-derived lineages in our ICM embryoids, as determined by pseudotime analysis, appears to be DVE/AVE. The establishment of the DVE/AVE and its confinement to a particular area relies on a dynamic equilibrium between NODAL and BMP signaling [92]. Respectively, these signaling pathways either facilitate [93–95] or constrain [60] the development of the DVE/AVE. *In vivo*, the ExEc plays a role in directly supporting BMP signaling and indirectly supporting NODAL signaling [95, 96]. Since our culture protocol contains serum, we believe that small variations in the serum that decrease NODAL and/or likely increase BMP signaling, can affect this dynamic equilibrium and sometimes result in the formation of cell population heterogeneity in the DVE/AVE-like cells which may differ from *in vivo* embryos. The low expression of NODAL antagonists (which are also effector genes of NODAL-signaling) in the DVE/AVE-like cells of our ICM embryoids for example, may be due to the rapid decrease of NODAL signaling in both the epiblast- and PE-derived lineages. The developmental trajectory of the PrE-like cells in our ICM embryoids nevertheless directs the emVE towards cells with a DVE/AVE-like profile.

Besides providing a robust non-integrated stem cell-based embryo model to study development; ICM embryoids offer valuable insights for improved ETX embryoid production. Elimination of i*Gata6* cells from the epiblast, as development proceeds, underscores that mouse stem cells in embryo models do not inadvertently assume each other’s functions. Although chimera experiments suggest lineage restriction in mouse early lineage stem cells [97–100], this restriction can thus be extrapolated to ES cells directed to become PrE. This is pivotal for evaluating the effects of mutations within specific given lineages on development. Furthermore, the smaller size of ICM embryoids, resembling the rosette stage morphologically [38], suggests that to better model post-implantation development, smaller numbers of ES cells should be favored when assembling i*Gata6* ES cells with ES cells not overexpressing *Gata6* to produce ETX embryoids. However, achieving a balance between ‘wild-type’ and i*Gata6* ES cells becomes more intricate with fewer cells. These findings emphasize the delicate equilibrium of lineage-restricted stem cell numbers in these experimental set-ups, which is pivotal yet very intricate when modeling early embryonic events.

In conclusion, the observed resemblance between *in vivo* epiblast development and *in vitro* ICM embryoids underscores its relevance for developmental biology research, validating decades of accumulated research within a single model. ICM embryoids offer insights into the specification of the PrE and the epiblast, the transition from naïve to primed pluripotency, and the anterior developmental fate of the epiblast in the absence of an ExEc. ICM embryoids not only offer an ethical alternative to traditional animal experimentation but could also facilitate studies on lineage interactions using mutated cell lines, thereby addressing questions that would be difficult or impractical to explore through conventional animal research.

## Materials and Methods

### Mouse ES cell culture

Mouse ES cells were derived and cultured as previously described [101, 102]. In summary, ES cell lines were generated and cultured on cell culture plates that had been pre-coated with 0.2% gelatin and irradiated mouse embryonic fibroblasts (MEFs, density 3×10^4^/cm^2^). The used ES cell lines comprised a male *Actin*-EGFP ES cell line (129S2/SvHsd-C57Bl/6JOlHsd) and a male ES cell line (129S2/SvHsd). ES cells were nurtured in ES cell medium containing DMEM (Gibco 41966-029), 15% fetal bovine serum (FBS), 100U/ml penicillin-streptomycin, 100µM non-essential amino acids (NEAA), 0.1 mM β-mercaptoethanol, and 20 ng/ml leukemia inhibitory factor (LIF). This medium was further supplemented with 2i, consisting of 1µM ERK inhibitor PD0325901 (Axon Medchem Cat 1408) and 3µM GSK3 inhibitor CHIR99021 (Axon Medchem Cat 1386). Cultures were maintained at 37°C under 5% CO_2_ atmosphere. All mouse ES cell lines were passaged using 0.25% trypsin-EDTA (5 to 7 min at 37°C) for dissociation of the cells. Trypsin-EDTA was subsequently inactivated with an equivalent volume of FBS, the cell suspension was then centrifuged at 1000rpm for 5min and suspended in their respective cell culture medium. Cells were split in ratios 1:6-1:24.

### Generation of Gata6 inducible ES cell lines

For the generation of *Gata6* inducible (i*Gata6*) ES cell lines an inducible dCas9-VPR system was used [103]. The inducible dCas9-VPR cassette was cloned into one allele of the safe harbor TIGRE locus along a constitutive M2rTTA cassette into the other TIGRE locus allele [104] of 129S2/SvHsd ES cell lines. Correct targeted ES cell lines with a normal karyotype were subsequently targeted with a PiggyBac vector containing a Cherry fluorescent reporter along two guides specific for the promoter of *Gata6* (GCTAGCCCTGTCCTCAGAGT and GGACACCAAGGGAGGGGAAG at -184bp and -238bp from the transcription start site, respectively) (Fig. S1A). Positive ES cell clones were picked, expanded and exposed to Doxycycline (2µg/ml) for 24h to assess their competence to upregulate *Gata6*.

### RNA isolation, cDNA synthesis and real-time PCR

Cells were collected and stored in lysis buffer (Promega Reliaprep RNA Cell Miniprep, cat z6012) at -20°C until further processed. Following thawing, RNA was isolated using the kit (Promega Reliaprep RNA Cell Miniprep, cat z6012) and eluted in 50µl water, all according to the manufacturer’s instructions. In total, 500-1000ng of RNA was reverse transcribed into cDNA using random hexamers and Superscript III reverse transcriptase (Thermo Fisher, cat no. 18080093) according the manufacturer’s protocol. Following dilution of the cDNA with water (1:5), expression levels were assessed via real-time PCR using a BioRad CFX384 Real-time PCR machine. The 10µl reaction mixture contained 2x SYBR Green PCR Master Mix (Promega, GoTaq qPCR Master, cat.no. A6002), 2 µM primer and 1 μl of diluted cDNA. After an initial hold at 95°C for 10 min, reaction mixtures underwent 40 cycles of 30 sec at 95°C and 1min at 60°C. The sequences of the used real-time primers can be found in Table 1 below.

**Table 1.**
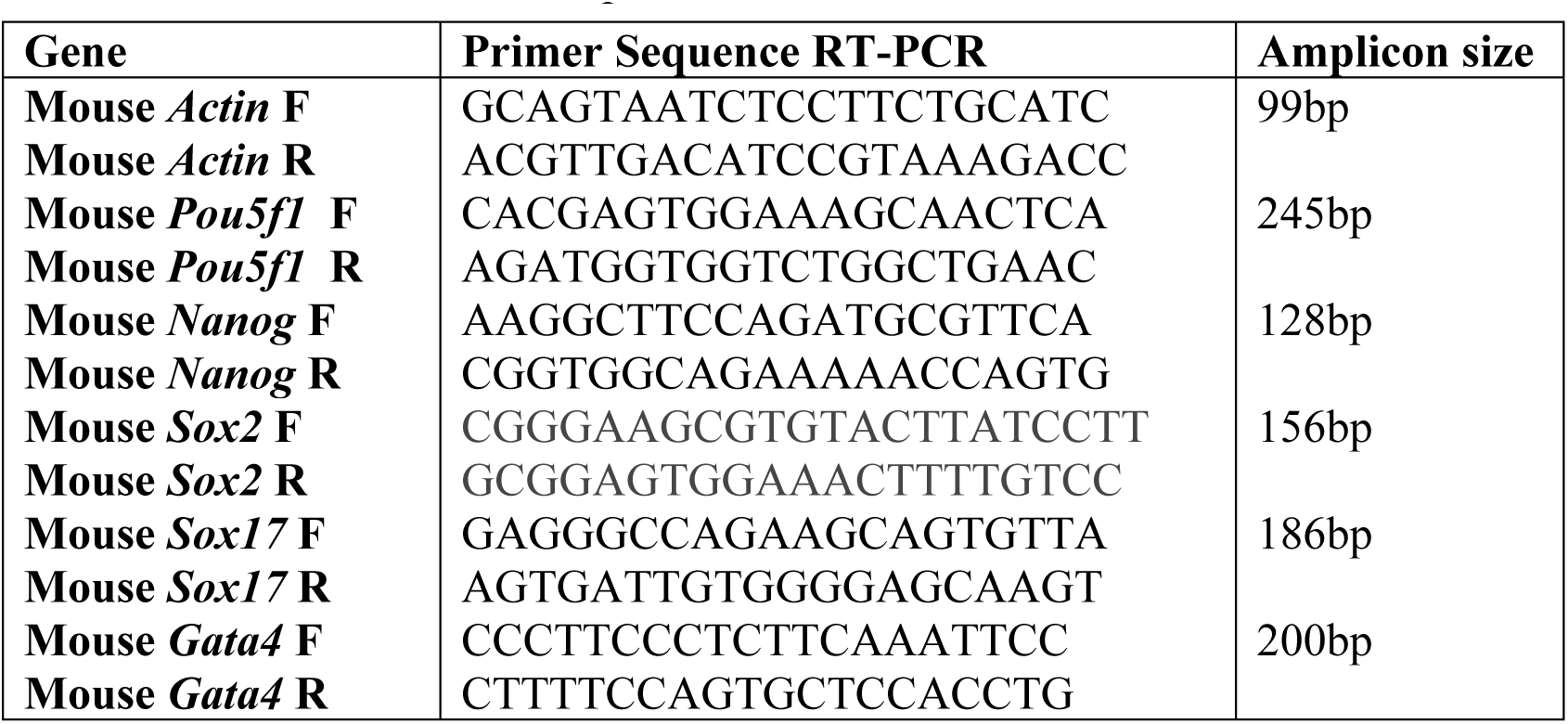
Used murine real-time primers.

### Aggrewell set-up,ES cell seeding and embryoid body/ICM embryoid culture

Six hours before trypsinization of the *Actin*-EGFP and i*Gata6* ES cells, i*Gata6* ES cells were exposed to Doxycycline (2µg/ml). Aggrewell 400µm plates were filled with 1ml of anti-adherence solution, centrifuged at 4000rpm for 5 min to remove air bubbles and incubated for 30min at room temperature. Following removal of the anti-adherence solution, wells were washed with 2ml PBS and filled with 500µl ETX medium (DMEM Gibco 41966-029, 15% FBS, 100µM NEAA, 1mM sodium pyruvate, 2 mM L-alanyl-L-glutamine, 100U/ml penicillin-streptomycin, 0.1mM β-mercaptoethanol and 20 ng/ml leukemia inhibitory factor (LIF)). ES cells were trypsinized using 0.25% trypsin-EDTA (5 to 7 min at 37°C). Following inactivation of the TE using ETX medium, cells were pipetted into single cells and twice washed with ETX medium. To mimic ICM dynamics, 24,000 to 30,000 cells diluted in 1ml were added per well (20 to 25 cells per µwell). The seeding day of the ES cells was considered as day 0. The Aggrewell plates were refreshed on day 2 with ETX medium devoid of LIF and on day 3 with IVC1 medium as previously described [7]. To assess the impact of Hippo pathway modulation on the positional differentiation of PrE, LPA sodium salt (500nM-5µM) or YAP-TEAD Inhibitor 1 (50nM -10µM) was added to the culture medium during the seeding of ES cells. Additionally, to investigate the effect of TGFβ-signaling inhibition on cellular arrangement, 10µM SB431542 was supplemented to the culture medium during the seeding process.

### Immunostaining

ICM embryoids were fixed in 4% paraformaldehyde for 20 min, washed in PBS (3×5 min), permeabilized with 0.25% Triton X-100 in PBS for 20min, washed in PBS (3×10 min) and incubated overnight in blocking solution (0.5% BSA, 1% Tween-20 in PBS) at 4°C (up to one week). Following blocking, ICM embryoids were exposed overnight to primary antibodies diluted in blocking solution at 4°C. After washing in blocking solution (3×10 min), ICM embryoids were incubated for an additional 2 days with the secondary antibody diluted in blocking base at 4°C. After 2 days, ICM embryoids were rinsed in PBS (3×10 min), Hoechst33342 20ug/ml was added to the first wash. ICM embryoids were embedded in Matrigel drops and imaged using a Leica SP5 Intravital microscope with a HCX APO 20x water-immersion objective. Images were processed with ImageJ. Used antibodies can be found in Table 2 below.

**Table 2.** Used antibodies for immunostaining.

| Target | Cat. Number | Dilution | Host |
| --- | --- | --- | --- |
| FOXA2 | Cell Signaling 8186T | 1:200 | Rabbit |
| GATA6 | R and D AF1700 | 1:500 | Goat |
| GATA4 | Santa Cruz sc9053 | 1:200 | Rabbit |
| POU5F1 | Santa Cruz sc5279 | 1:500 | Mouse |
| OTX2 | R and D AF1979 | 1:100 | Goat |
| TBR2 | Abcam AB23345 | 1:200 | Rabbit |
| POU3F1 | Absea 060204E0 | 1:50 | Mouse |
| Phalloidin-546 | Thermofisher | 1:200 | - |
| Mouse Alexa 633 | A21146 | 1:500 | Donkey |
| Rabbit Alexa 488 | A21206 | 1:500 | Donkey |
| Goat Alexa 555 | A21432 | 1:500 | Donkey |

### Processing of ICM embryoids for 10X Genomics

Naïve ES cells (*Actin*-EGFP and i*Gata6* ES cells) and ICM embryoids were collected on day 1, day 2, day 3 and day 4. Per day, 100-300 ICM embryoids were processed for single-cell RNA sequencing using the 10X Genomics Chromium Platform as previously described [44]. Briefly, ICM embryoids were enzymatically digested with TrypLE Express dissociation reagent (Life Technologies) for 5 min at 37 °C followed by gentle mechanical dissociation using small pipette tips. After inactivation with an equivalent volume of FBS, the single cells were filtered through a Flowmi Tip Strainer with 40 µm porosity (ThermoFisher Scientific, cat.no.136800040), washed and resuspended in PBS with 0.4% BSA.

### Single cell library preparation and sequencing

Single cells were processed on the 10X Genomics Chromium Platform using the Chromium Next GEM Single Cell 3′ Reagent kit v3.1 with dual indexing (10X Genomics) following the manufacturer’s protocol. Briefly, 15,000 cells were loaded in each channel of a chip to be partitioned into gel beads in emulsion (GEMs). Within GEMs, cells were lysed followed by barcoded reverse transcription of RNA. Breaking of the GEMs was followed by amplification, fragmentation and addition of adapter and sample index. Libraries were pooled and sequenced 28-10-10-90 cycles on an Illumina NovaSeq6000 instrument aiming for a minimum coverage of 25,000 raw reads per cell.

### Pre-processing of scRNA-seq data

10× Genomics Cell Ranger mkref v6.0.0 [105] was used to create a custom reference package containing the mm10 reference genome and the sequences of EGFP and Cherry. The raw FASTQ files of each day were processed separately using cellranger count with the custom reference. The counts were preprocessed using R v4.0.5 and Seurat v4.3.0 [106]. SoupX v1.5.2 [107] was used to remove ambient background RNAs from the datasets. We removed doublets using doubletfinder v2.0.3 [108] with the settings pN = 0.25, pK = 0.09, nExp = 0.05 * number of cells, PCs = 1:10. Cells with less 2,000 or more than 8,000 detected genes, more than 70,000 counts or more than 10 percent of mitochrondrial reads were removed. Genes expressed in less than 3 cells were removed. GFP and *Cherry* were removed from the matrix and their counts were added as separate meta columns. Cells without EGFP and *Cherry* expression were removed from the dataset. Cells with at least one *Cherry* read were labeled as *Cherry* positive cells, while the remainder were labeled as EGFP positive cells.

### Integration of scRNA-seq datasets

Integration of the cells of the different days were integrated using scanpy v1.9.1 [109]. To allow an equal representation of the different days, we randomly downsampled the cells from each dataset to the lowest number of cells per dataset (i.e. 2064 cells). The cells were integrated using the integrate_scanpy function of scanorama v1.7.1 [110] based on the expression of variable genes from the highly_variable genes function (min_mean=0.0125, max_mean=3, min_disp=0.5, batch_key=“day”) and filtering for genes that are variable in at least two of the days. Based on the scanorama embedding, an Uniform Manifold Approximation and Projection (UMAP, [111]) was created using the scanpy v1.9.1 neighbors (n_pcs=30, use_rep=”Scanorama”) and umap functions. The cells were clustered using the Louvain algorithm [112] with resolution=1. Cluster 7 containing 21 cells was removed from the dataset, as these cells represent differentiated cells in the Naive state. Based on the proportion of EGFP positive and Cherry positive cells, cluster 0, 1 and 2 were annotated as EGFP clusters, while cluster 4, 5 and 6 represent Cherry clusters and cluster 3 consists of Naive cells. Expression of lists of marker genes was plotted using scanpy dot plot with standard_scale=“group”. Expression of separate genes was also visualized on the UMAP using seurat FeaturePlot. The EGFP positive lineage cells were categorized into three pluripotency states: naive, primed, or an intermediate state based on the expression of *Klf4, Zfp42, Pou3f1*, and *Otx2*. Cells expressing either *Klf4* or *Zfp42* were labeled as naive, while those expressing *Pou3f1* or *Otx2* were labeled as primed. Cells expressing both naive and primed markers were classified as intermediate (naive-primed), whereas cells without any label were excluded. The distribution of pluripotency labels per day was visualized in a stacked bar plot.

### Pseudotime analysis

Monocle3 v1.0.0 [113] was used to perform pseudotime analysis. The cells were preprocessed using the preprocess_cds function with num_dim=100 and aligned using align_cds with alignment_group=“day”. Learn_graph was run with use_partition=T based on the clusters identified by the Louvain algorithm. Finally, the cells were ordered using order_cells with reduction_method=UMAP and the cells at the tip of cluster 3 as root_cells. The pseudotime of the cells was visualized on the UMAP generated by scanorama. We created a heatmap with the expression of marker genes along the pseudotime by ordering the cells of the Cherry clusters by pseudotime, smoothing the expression values using the R stats function smooth.spline function with df=3 and converting these smoothed values to z-scores.

### RNA velocity analysis

Velocyto v0.17.17 [114] run10x was run on each day seperately to create loom files with spliced and unspliced counts. The loom files were analyzed using scvelo v0.3.0 [115]. The counts were pre-processed using scvelo filter_and_normalize with 20 min_shared_counts and 2000 n_top_genes, scanpy pca and scanpy neighbors with 30 n_pcs and 30 n_neighbors. We ran scvelo moments and recover_dynamics to compute moments for velocity estimation and recover the full splicing kinetics, respectively. Finally, scvelo velocity and velocity_graph was run with the mode set to “dynamical” to obtain the velocity estimates. The velocity of the cells was plotted on the scanorama UMAP using scvelo velocity_embedding_stream with smooth=0.8 and min_mass=1.

### Differential expression analysis

We performed differential expression analysis between the Cherry positive cells on day 1 and 2 and the Naive cells to uncover which genes are activated during development. Differentially expressed genes (DEGs) were identified using the seurat FindMarkers function with min.pct=0.1 and logfc.threshold=0.1 and filtered for upregulated genes. The top 50 DEGs were subjected to pathway analysis using EnrichR v3.2 [116] with the Reactome 2022 database [117]. The most significant pathways were visualized in a bar graph, displaying the -log10(P-value) and the number of DEGs associated with each pathway. We then focused on pathways with the highest number of DEGs (excluding child terms) and assessed the enrichment of their corresponding DEGs using the ‘score_genes’ function from scanpy (with a control group of 500 genes), which results were plotted on UMAPs.

### CellphoneDB

Cell-cell interactions between the different groups of cells (i.e. cells per day per lineage) were identified using CellphoneDB v5.0 [61], with the CellphoneDB v5.0 interaction database. Mouse genes were converted to their respective one-to-one human orthologue using the Ensembl v98 database. The statistical method of CellphoneDB (cpdb_statistical_analysis_method) was run, focusing exclusively on interactions within each day, as specified in a microenvironment input file. The number of significant interactions (p-value < 0.05) for each possible pair was counted and visualized in a heatmap. The different interaction combinations were split according to the lineage from which the interaction originates. Active interactions for each lineage were selected based on the condition that they were significant on at least one day. Within each lineage, active interactions were annotated by pathway, and pathways with at least three active interactions (i.e. active pathways) were visualized in dot plots. The number of active interactions originating from each lineage was split by pathway (limited to pathways active in at least one lineage) and plotted in a barplot.

### Pathway analysis

Based on literature research we selected known effectors, ligands, and receptors for several pathways including JAK/STAT, WNT, ACTIVIN/NODAL, MAPK, Hedgehog, NOTCH and BMP. The expression of these known genes per day of each lineage was visualized using dot plots. For each pathway, we calculated the average expression per day per lineage and transformed these values to z-scores. For each lineage separately, these scores were visualized in a diagram over time, where the differences between days were illustrated as a gradient.

### Comparison of cell types between ETX and ICM embryoid data

The scRNA-seq data from ETX embryoids was pre-processed as previously described [7]. The percentage of cell types present in ETX and ICM embryoid data were assessed by selecting cells with a combination of marker genes expressed based on minimum CPM thresholds. Primitive Streak cells were selected based the expression of both *Fgf8* and *Fst*. Percentages of mesoderm and endoderm cells was based on the expression of *T* and *Foxa2*, respectively.

### Comparison to in vivo scRNA-seq

An integrated scRNA-seq dataset of cells from embryos, blastoids and stem cells was downloaded from Posfai et al., 2021 (GSE145609) [39]. The original UMAP and cell type labels were added to the scanpy object and all *in vitro* cell clusters and clusters with less than 15 cells were removed, resulting in a UMAP containing only embryo cells. Genes specific to Naive cells, GFP positive cells per day and Cherry positive cells per day were identified through a differential expression analysis using scanpy rank_genes_groups with a t-test between these groups. For each ICM embryoid group, the top 500 differentially expressed genes (DEGs) were selected as group markers. Subsequently, the enrichment of these marker genes in the *in vivo* cells was calculated using scanpy score_genes with a random reference gene list size of 500. The enrichment in each cell was visualized on the UMAP. Moreover, the top 50 cells with the highest enrichment were selected and highlighted on the UMAP showing the most similar cells for each ICM embryoid group. For each top cell, the *in vivo* cell type annotation was extracted and plotted in a stacked barplot showing the distribution of different cell types across ICM embryoid groups. Cell types observed in at least 10 top cells across all groups were individually labeled, while less frequently found cell types were classified together as “Other”.

Moreover, scRNA-seq data from Nowotschin et al., 2019 (GSE123046) was re-analyzed [48] Count tables from E3.5 to E7.5 were downloaded and merged into a scanpy object. Low-quality cells were removed using automatic thresholding based on median absolute deviations (MAD), as described in Germain et al., 2020 [118]. Cells were flagged as outliers if they deviated more than 5 MADs in log(total_counts+1), log(n_genes_by_counts+1) or pct_counts_in_top_20_genes, and were subsequently removed. Cell type labels from the original study were added to the object, and filtered for cells annotated as ICM, epiblast, primitive endoderm, visceral endoderm, extra-embryonic visceral endoderm and embryonic visceral endoderm. The remaining cells were integrated using Harmony v0.1.5 [48]. Counts were normalized, the top 2,500 highly variable genes were selected, and the data were log-transformed using the normalize_counts, hvg_genes and log_transform functions, respectively. An augmented affinity matrix was constructed using sample time point information with the augmented_affinity_matrix function and used to generate a force-directed layout for visualization using the force_directed_layout function. As described above, the enrichment of the top 500 DEGs for each ICM embryoid group was calculated using scanpy score_genes. The top 100 cells with the highest enrichment for each group were then selected and collectively highlighted on the Harmony force-directed layout.

## Funding

This work has become possible through a ZonMW-PSIDER grant (10250022120002) awarded to JG.

## Author contributions

Conceptualization: CD

Methodology: CD, BT, OS

Investigation: CD, BT, OS, EB

Visualization: CD

Supervision: CD

Writing-original draft: CD

Writing-review and editing: CD, BT, OS, SM, DH, JG

## Competing interests

The authors declare no competing interests.

## Data and materials availability

All raw and processed scRNA-seq data generated in this study have been submitted to the NCBI Gene Expression Omnibus (GEO) under accession number <u>GSE261620</u>. Reused data were retrieved from GEO (<u>GSE145609</u> and <u>GSE123046</u>).

---

**Figure S1.**
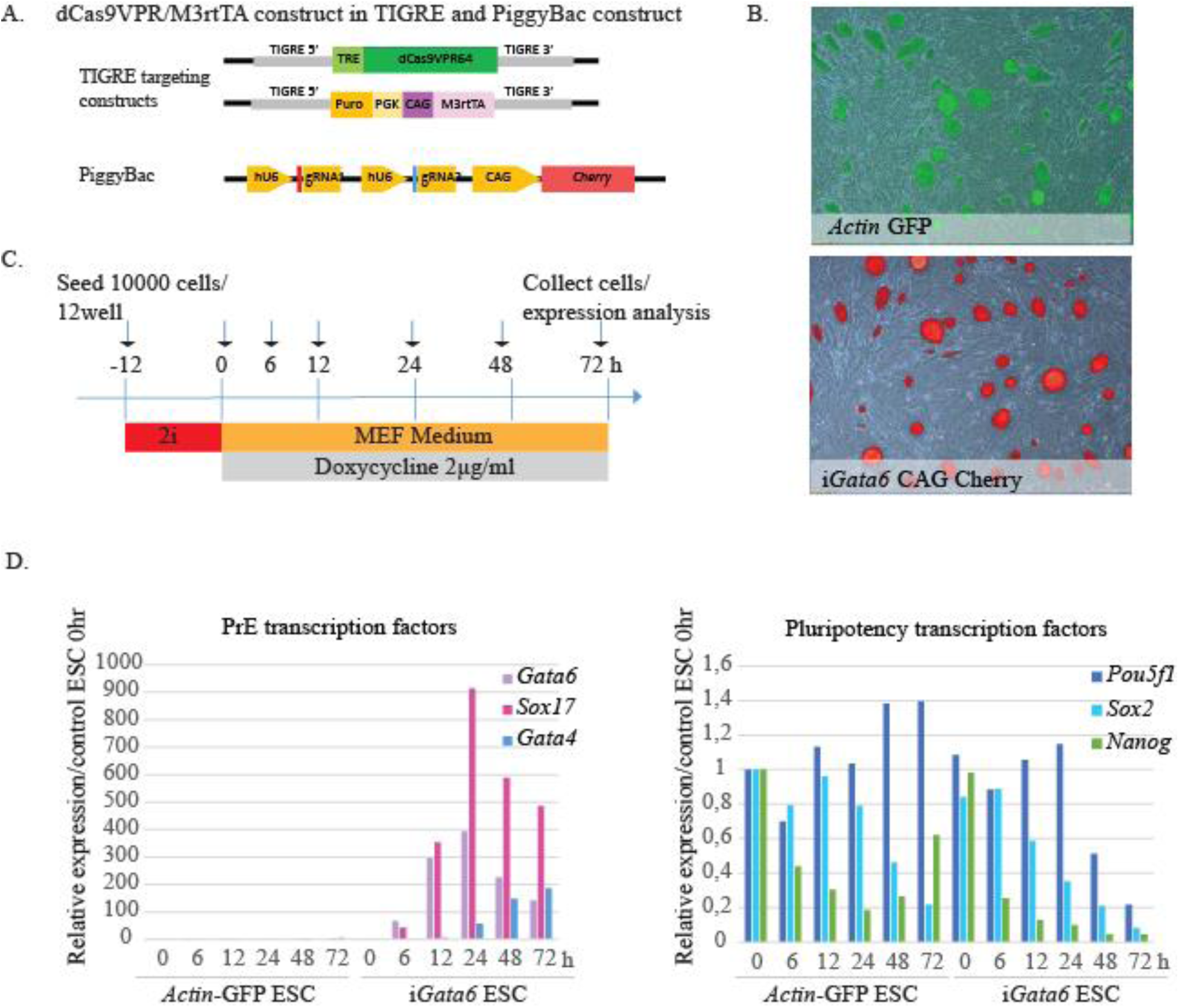
Construction and characterization of i*Gata6* ES cell lines. A. Schematic representation of DNA constructs to create i*Gata6* ES cell lines. Constructs with dCas9VPR64 and CAG-M3rtTA used for targeting the TIGRE locus (top) and a PiggyBac construct containing two different guide RNAs in tandem specific to the *Gata6* promoter (for details, see Materials and Methods) B. Photographs of cultured, undifferentiated “wild-type” (*Actin-* EGFP) and i*Gata6* ES cell lines. C. Procedure to test the temporal expression of genes encoding markers of PrE and pluripotency (i.e. transcription factors) during an induction experiment using doxycycline (see Materials and Methods). D. Steady-state mRNA expression profiles quantified using QPCR of PrE and pluripotency transcription factors in “wild-type” *Actin*-EGFP ES cells and i*Gata6* ES cells exposed to 2µg doxycycline/ml for 6, 12, 24, 48 and 72h, respectively, relative to 2i-cultured *Actin*-EGFP ES cell lines.

**Figure S2.**
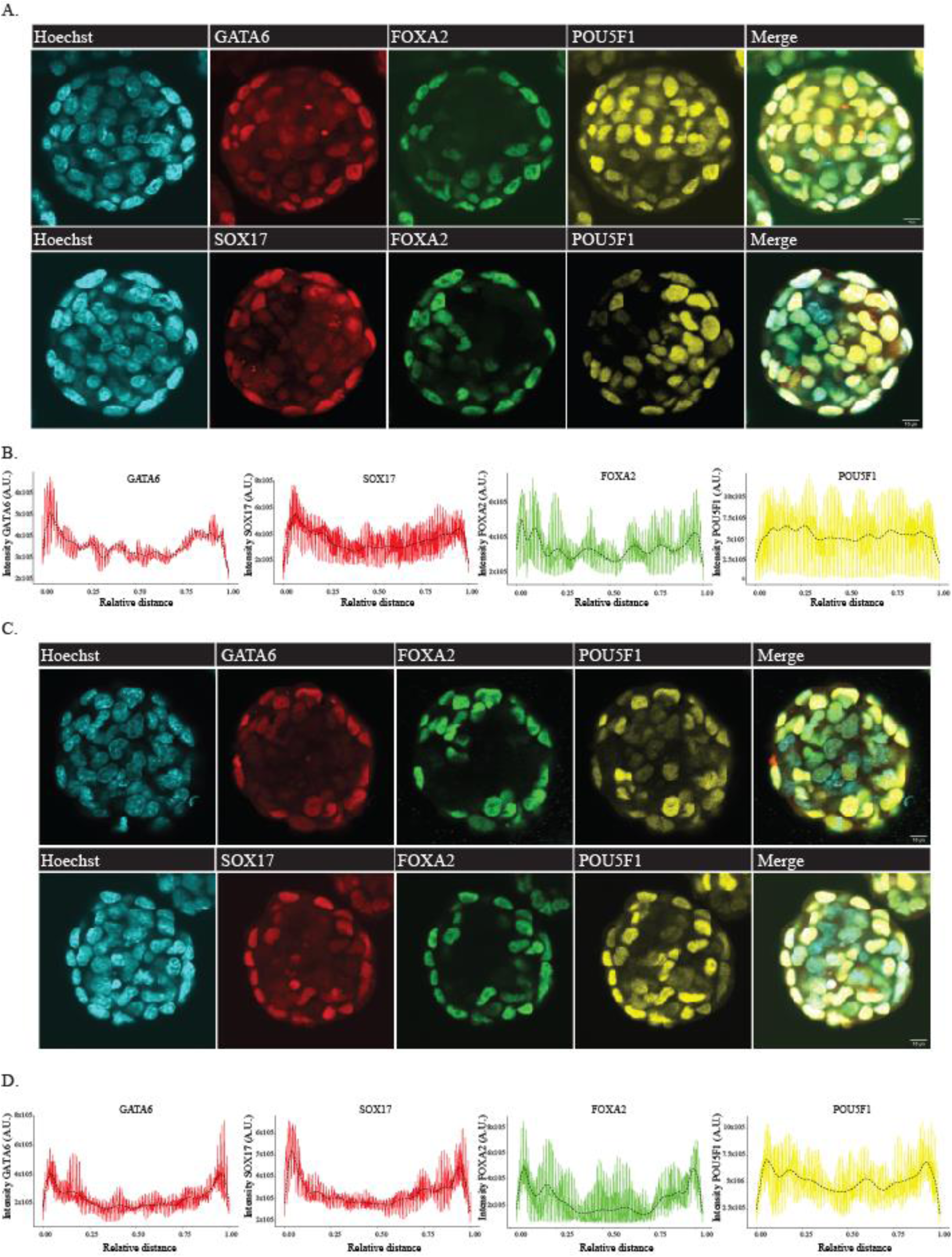
Position-dependent development of PrE in i*Gata6* embryoids. A. i*Gata6* EBs were collected on day 2 post-aggregation (following 6h Dox exposure) and assessed for PrE/VE and epiblast markers. In the top panel, i*Gata6* EBs stained for GATA6, FOXA2 and POU5F1. In the bottom panel i*Gata6* EBs stained for SOX17, FOXA2 and POU5F1, scale bar 10µm. B. Quantification GATA6, SOX17, FOXA2 and POU5F1 intensity profiles measured across multiple line scans (*n*=2-4) per EB (*n*=10 per staining). Trend analysis is performed using orthogonal polynomial fitting (dashed line). C. i*Gata6* EBs were collected on day 2 post-aggregation (following 30 h Dox exposure) and assessed for PrE/VE and epiblast markers. In the top panel, i*Gata6* EBs stained for GATA6, FOXA2 and POU5F1. In the bottom panels i*Gata6* EB stained for SOX17, FOXA2 and POU5F1, scale bar 10µm. D. Quantification of GATA6, SOX17, FOXA2 and POU5F1 intensity profiles measured across multiple line scans (*n*=2-4) per EB (*n*=10 per staining). Trend analysis is performed using orthogonal polynomial fitting (dashed line).

**Figure S3.**
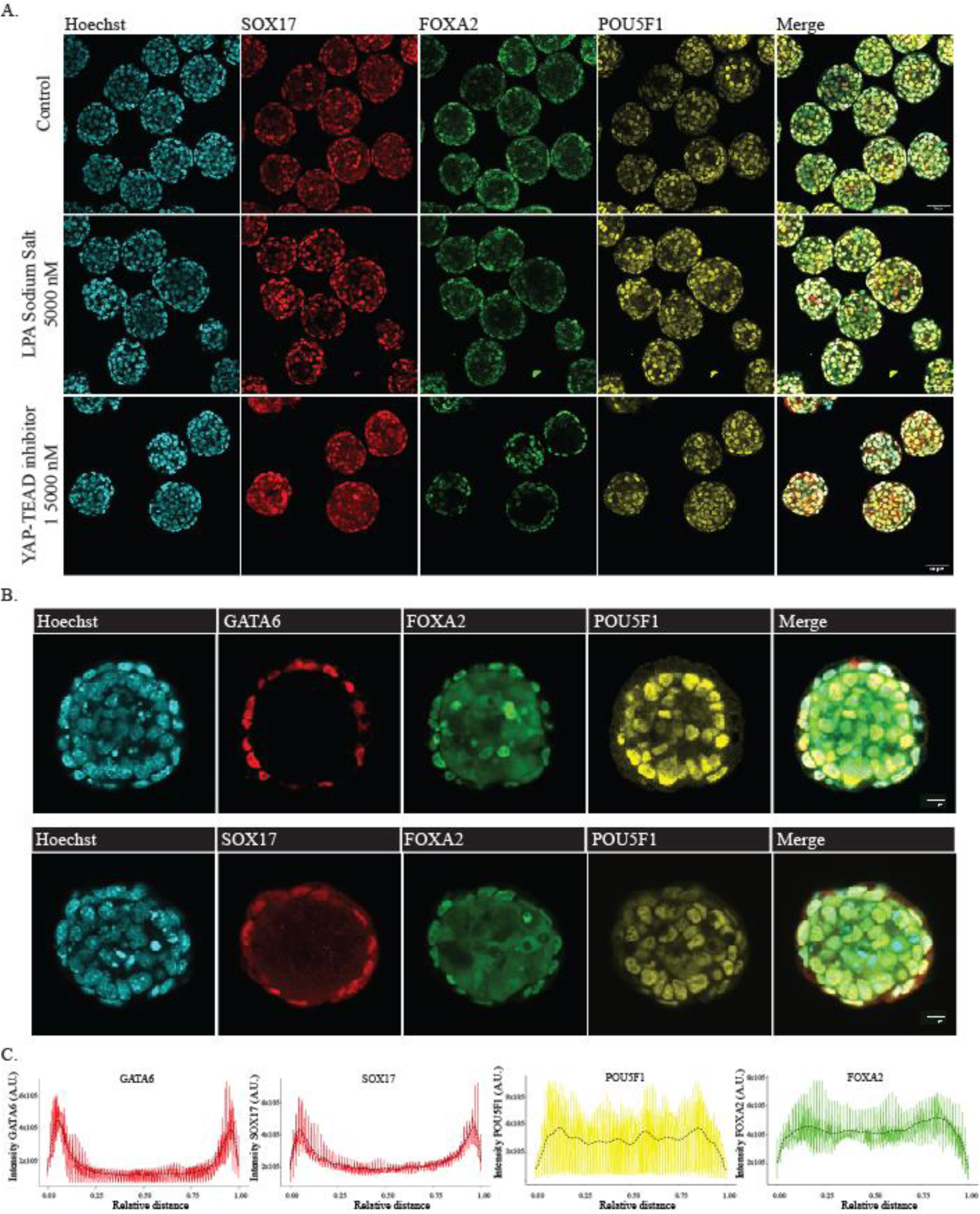
Position-dependent Hippo-independent PrE development and cell sorting facilitate PrE at the periphery of both i*Gata6* EBs and ICM embryoids. A. Assessment of influence of small molecules affecting Hippo pathway activity on position-dependent development of the PrE. i*Gata6* EBs collected on day 2 post-aggregation (following 6h of doxycycline exposure) and assessed for PrE/VE and epiblast markers; top panel, control; mid panel, 5µM LPA Sodium Salt and bottom panel, 5µM YAP-TEAD Inhibitor1. B. ICM embryoids were collected on day 2 post-aggregation (following 6h of doxycycline exposure) and assessed for PrE/VE and epiblast markers. In the top panels, ICM embryoids stained for GATA6, FOXA2 and POU5F1. In the bottom panels ICM embryoids stained for SOX17, FOXA2 and POU5F1, scale bar 10µm. C. Quantification of GATA6, SOX17, FOXA2 and POU5F1 intensity profiles measured across multiple line scans (*n*=2-4) per ICM embryoid (*n*=10 per staining). Trend analysis is performed using orthogonal polynomial fitting (dashed line).

**Figure S4.**
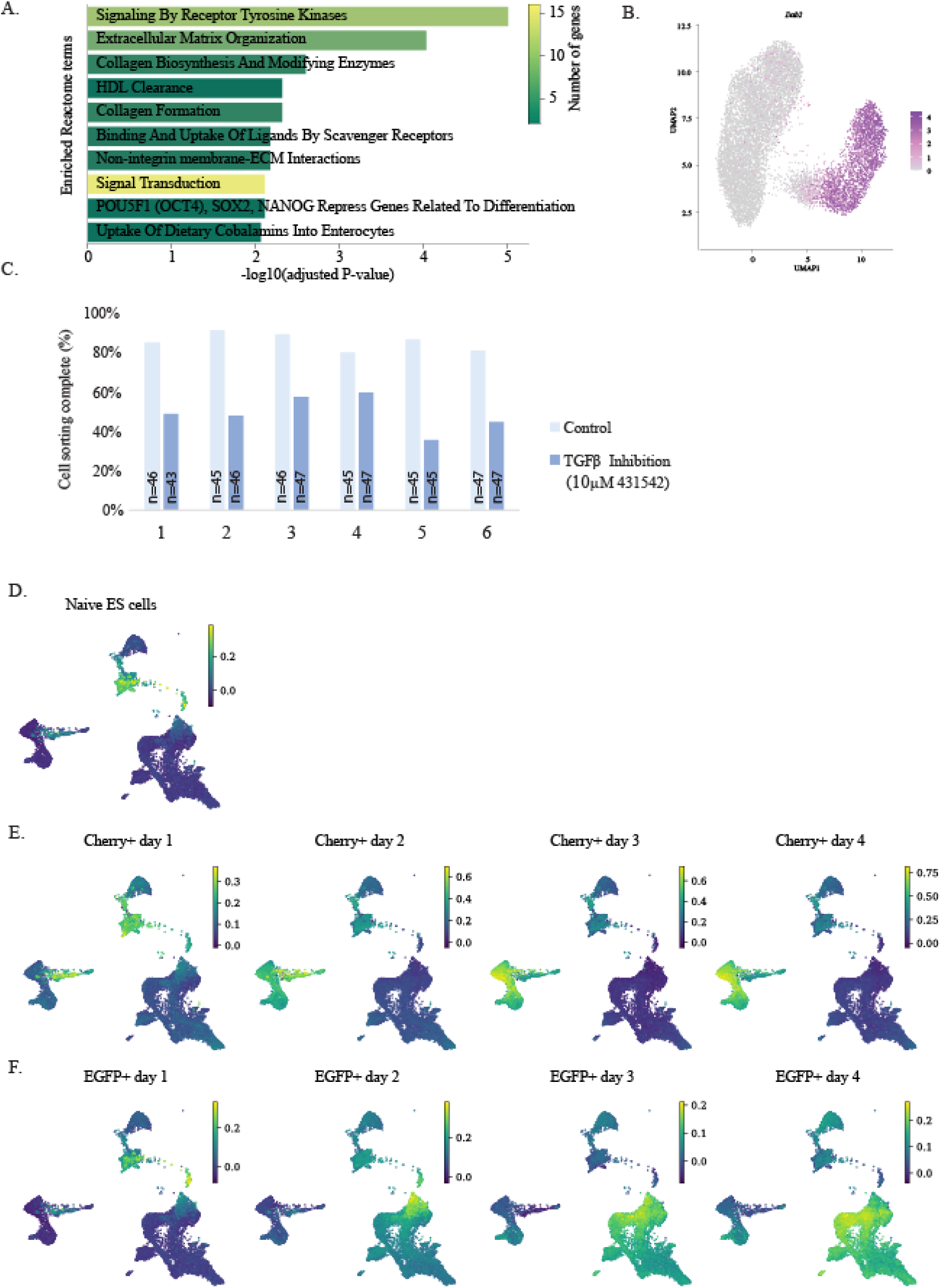
Exploring cell sorting mechanisms and developmental progression in ICM embryoids. A. Barplot showing the enriched Reactome pathways based on the top 50 upregulated genes in Cherry-positive cells of day 1-2 ICM embryoids compared to naïve ES cells. The bars are colored based on the number of upregulated genes associated with each pathway. B. Integration of scRNA-seq data of naïve ES cells and ICM embryoids (day 1-4). UMAP plot shows the normalized expression of *Dab2*. C. ICM embryoids composed of wild-type (*Actin*-EGFP) and i*Gata6* ES cells display perturbed, inefficient cell sorting following exposure to 10µM SB431542 resulting in TGFβ family inhibition. Bar plot depicting the average percentage of day 2 ICM embryoids displaying complete cell sorting, based on 6 biological replicates per condition. Total assessed ICM embryoids per replicate is displayed in bar chart. D. UMAPs showing enrichment of DEGs from each ICM embryoid cell cluster for each cell of the scRNA-seq dataset from [34]. E. UMAP is colored based on the enrichment of DEGs from naïve cells, F. Cherry positive cells in day 1 to 4 ICM embryoids, and C. EGFP positive cells in day 1 to 4 ICM embryoids.

**Figure S5.**
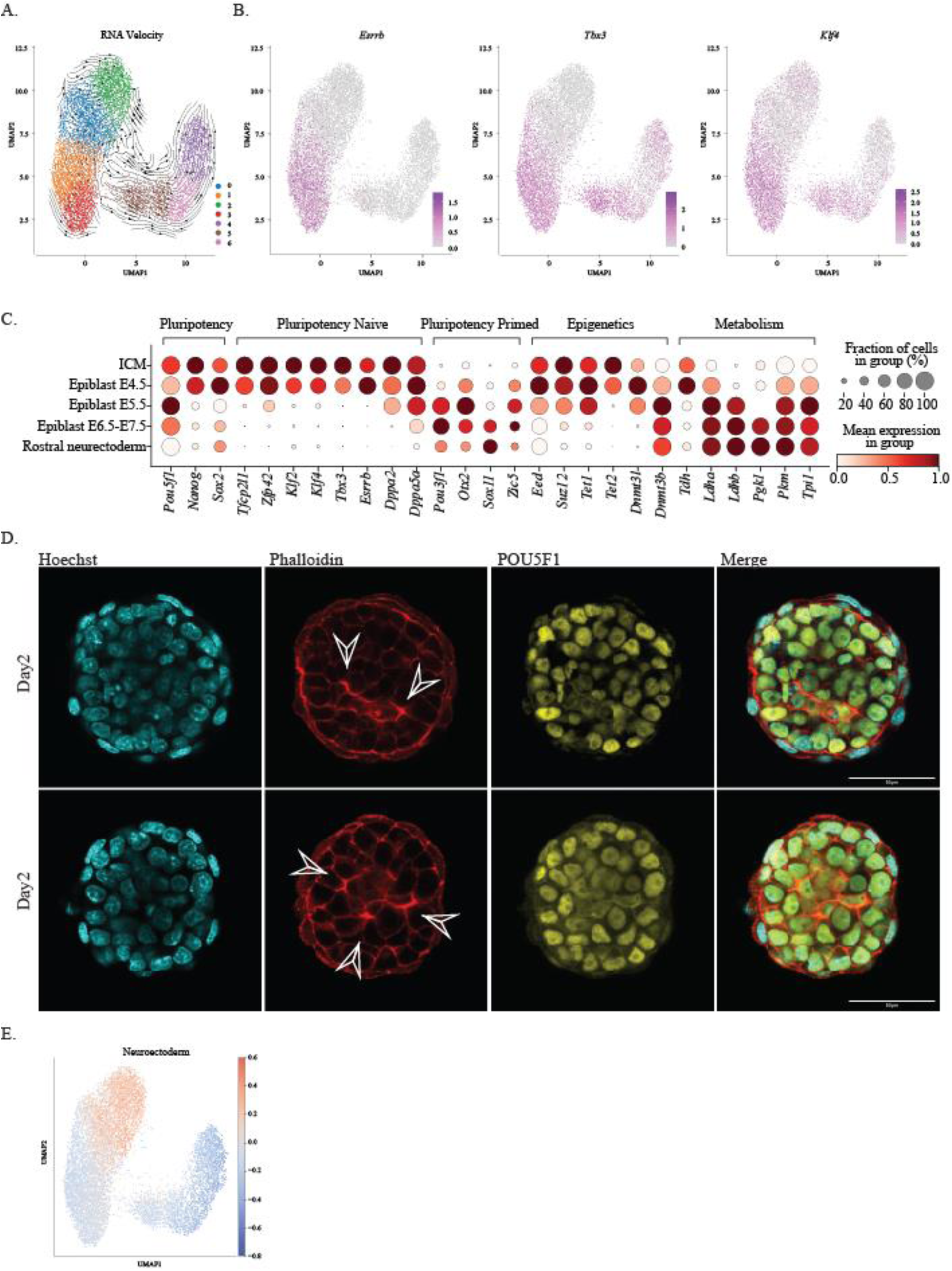
Naive to primed pluripotency transition occurs between day 1 and day 2 in epiblast-like cells of ICM embryoids. A. UMAP of integration of scRNA-seq data of naïve ES cells and ICM embryoids (day 1-4) colored by identified clusters with arrows indicating RNA velocity B. UMAP with normalized expression of *Esrrb, Tbx3* and *Klf4*. C. Dot plot showing expression of naïve and primed markers in EGFP-positive populations over time in correlated *in vivo* epiblast-derived tissues grouped by pathway. Color indicates normalized gene expression, while dot size reflects the percentage of cells expressing the gene. D. Lumen formation in day 2 ICM embryoids stained for Phalloidin and POU5F1 (arrows indicate lumen formation), scale bar 10µm. E. UMAP of integration of scRNA-seq data of naïve ES cells and ICM embryoids (day 1-4) showing rostral neuroectoderm signature based on the top 500 DEGs of *in vivo* rostral neuroectoderm cells.

**Figure S6.**
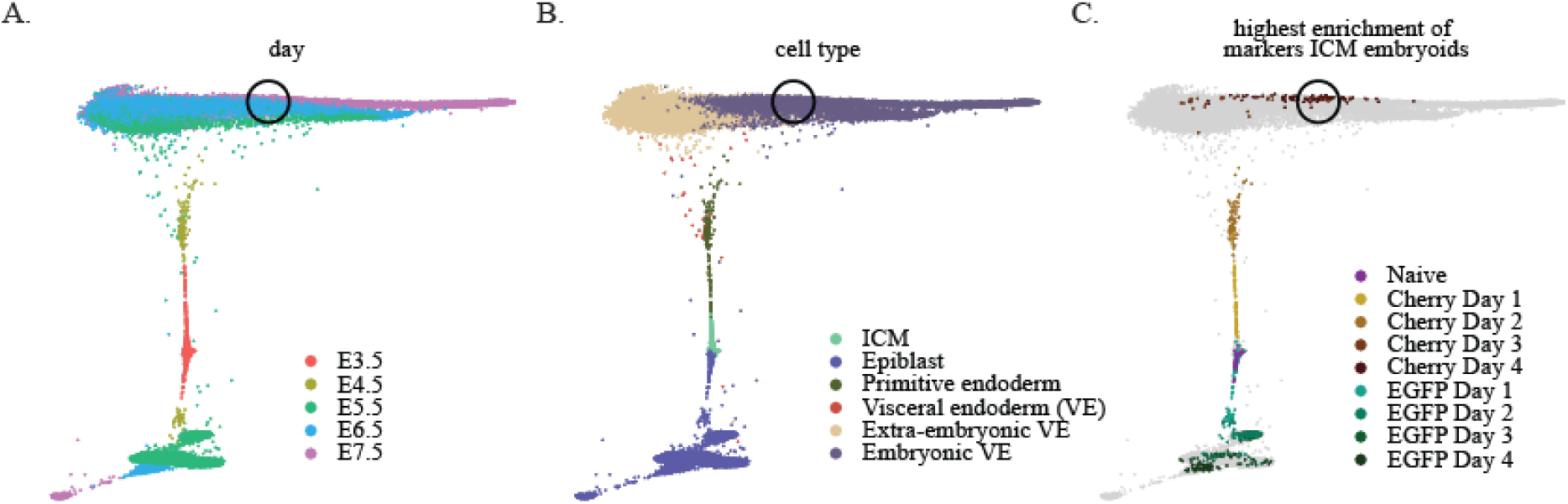
PrE-derived lineages resemble embryonic VE rather than extra-embryonic VE. A-B. Harmony force-directed layout of *in vivo* embryo cells from E3.5 to E7.5 [48], colored by day (A) and by cell type (B). ICM, inner cell mass. C. Harmony force-directed layout highlighting *in vivo* cells most similar to those from ICM embryoids. For each group of ICM embryoid cells over time (Naive, EGFP-positive, or Cherry-positive), the 100 *in* vivo embryo cells with the highest enrichment for the respective markers were identified and labeled. In panels A-C, a circle was added to indicate the region with the highest number of cells enriched for Cherry-positive day 4 markers.

**Figure S7.**
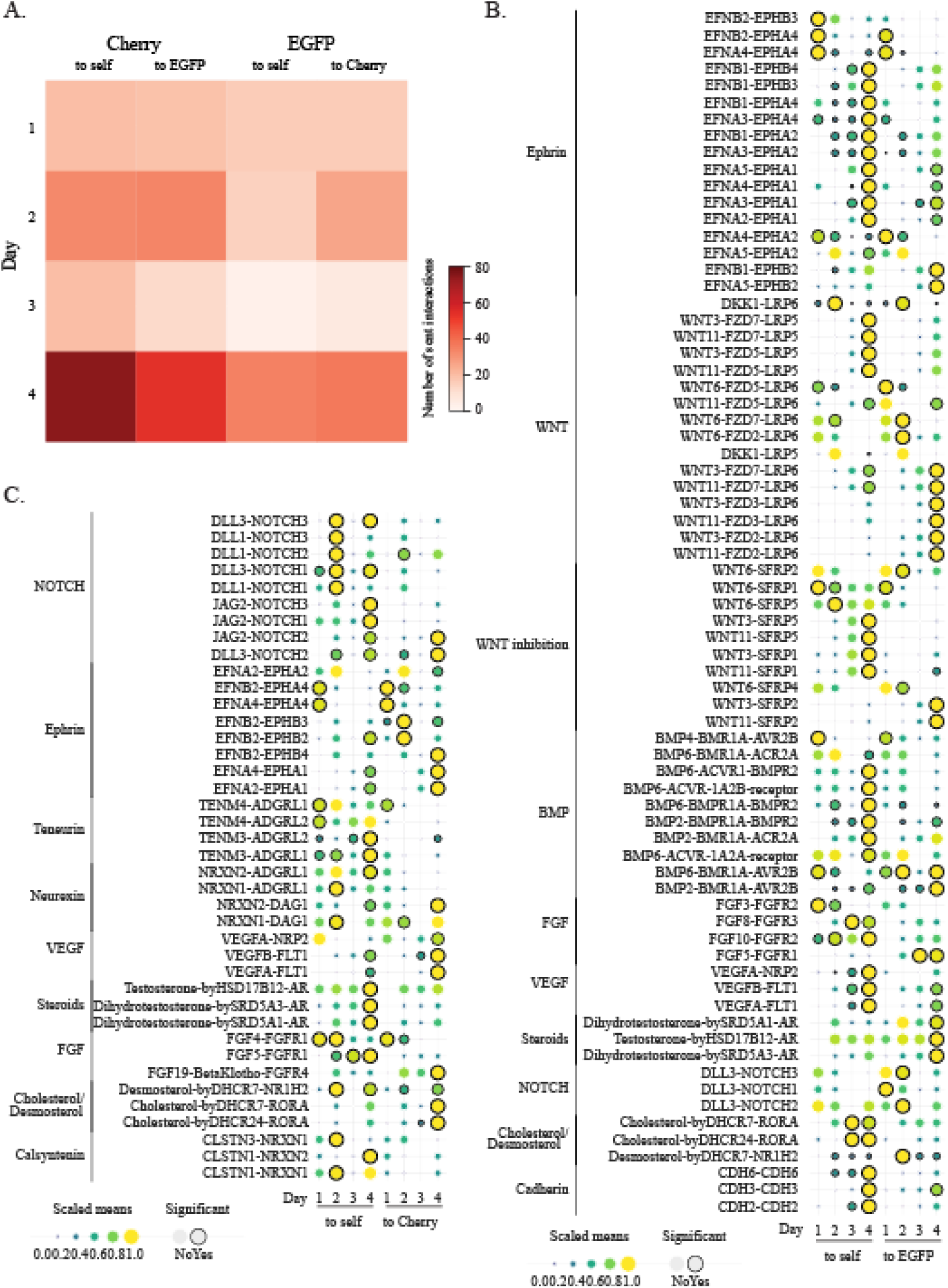
Cell lineage interactions detected by CellphoneDB. A. Heatmap showing the number of significant sent interactions identified by CellphoneDB between each cell type per day. B-C. Dot plot showing the CellphoneDB interactions significant in at least one combination within the EGFP (B) or Cherry (C) lineage, ordered by pathway. Only pathways with at least 3 significant interactions in the lineage are shown. Dot size and color indicate the mean interaction score and significant dots are highlighted with a black border.

**Figure S8.**
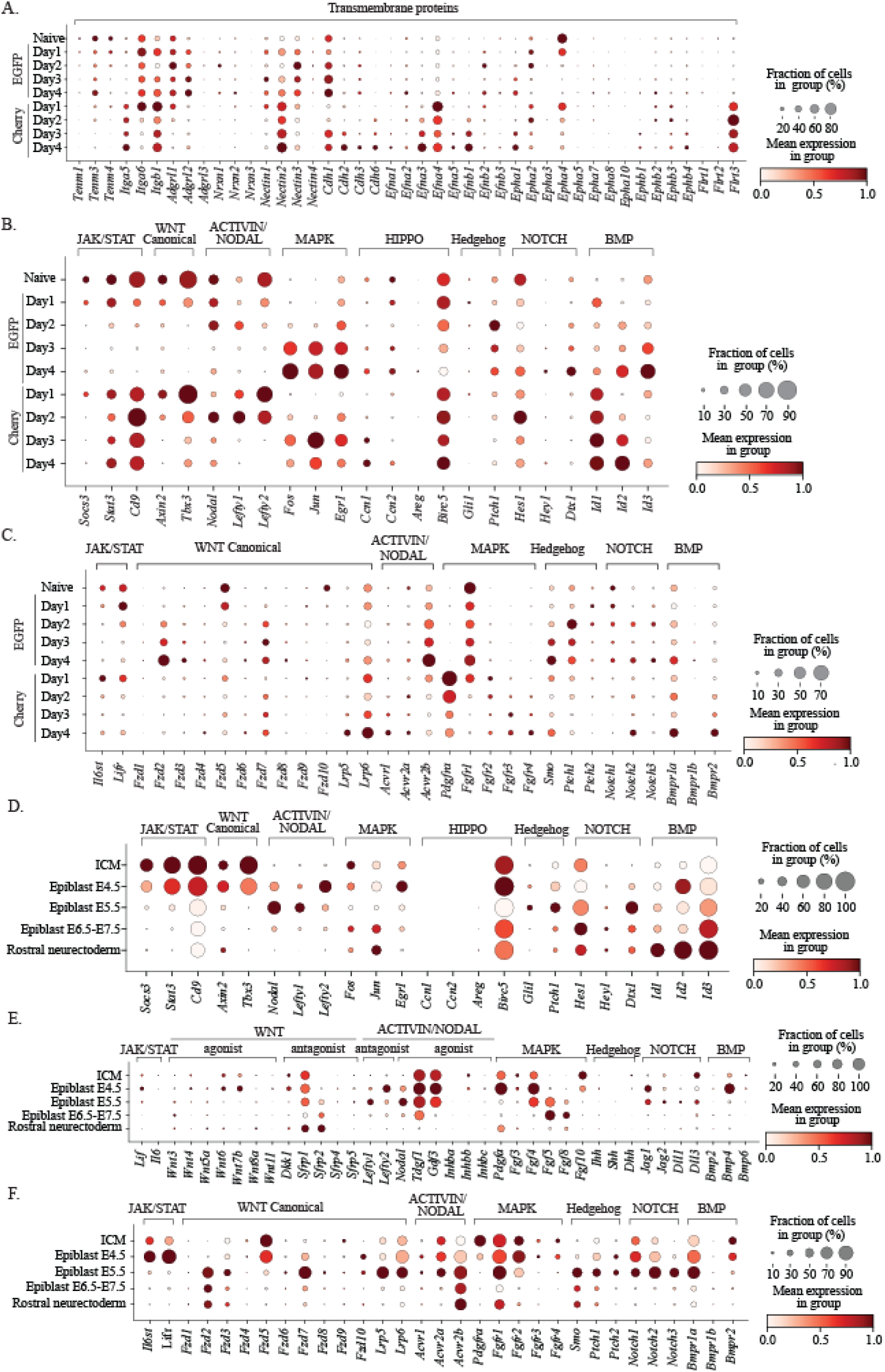
Normalized expression of genes in ICM embryoids and *in vivo* embryo culsters. A. Dot plot showing normalized expression of transmembrane genes in ICM embryoids grouped by pathway. Color indicates normalized gene expression, while dot size reflects the percentage of cells expressing the gene. B. Dot plot showing normalized expression of effector gene in ICM embryoids grouped by pathways. C. Dot plot showing normalized expression of receptors in ICM embryoids grouped by pathways. D. Dot plot showing normalized expression of effector genes for several pathways in embryo clusters shown in Figure 3C. Color indicates normalized gene expression, while dot size reflects the percentage of cells expressing the gene. E. Dot plot showing normalized expression of ligands for several pathways in embryo clusters shown in Figure 3C. F. Dot plot showing normalized expression of receptor genes for several pathways in embryo clusters shown in Figure 3C.

## Table legends

**Table S1.**
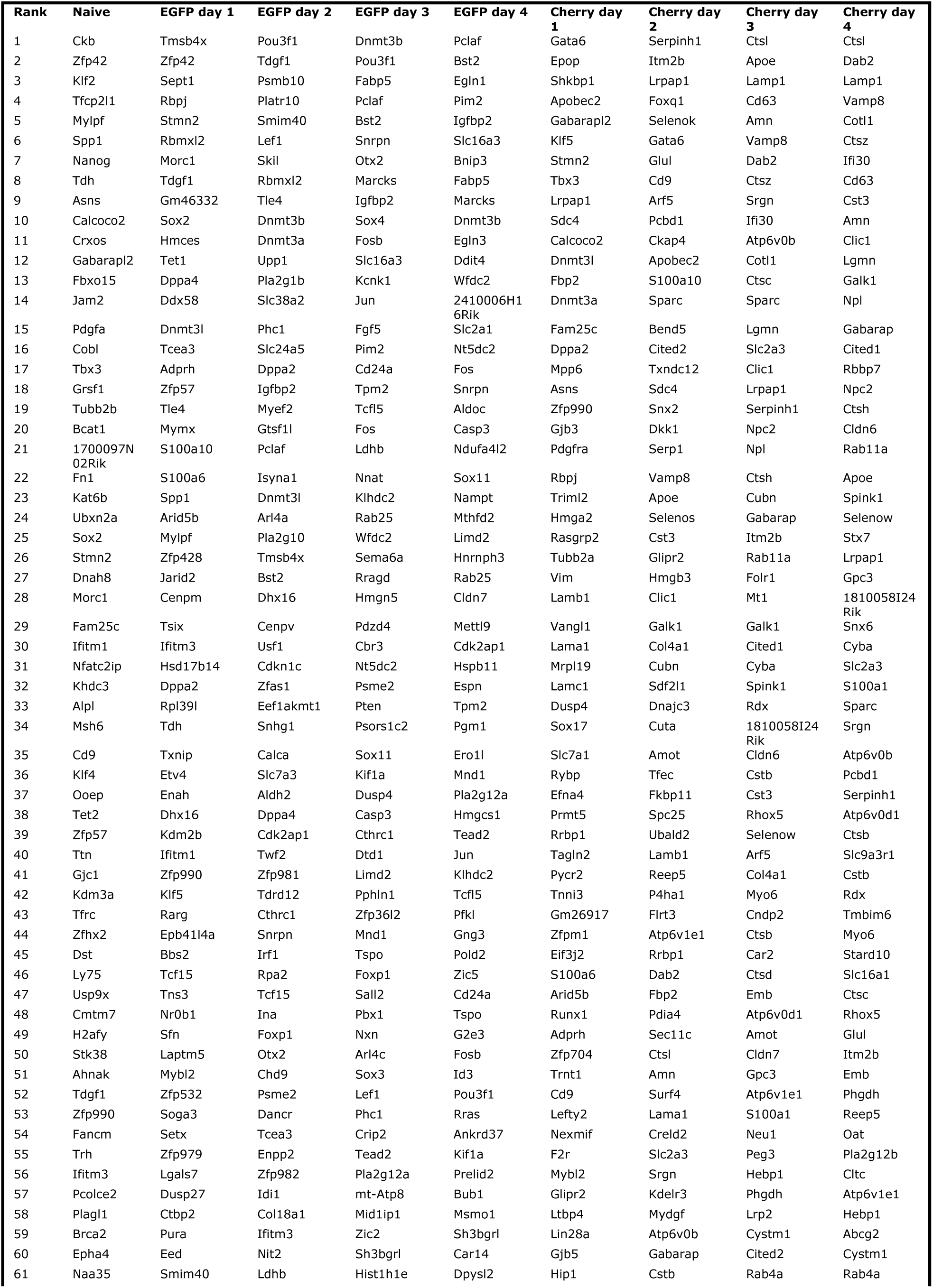
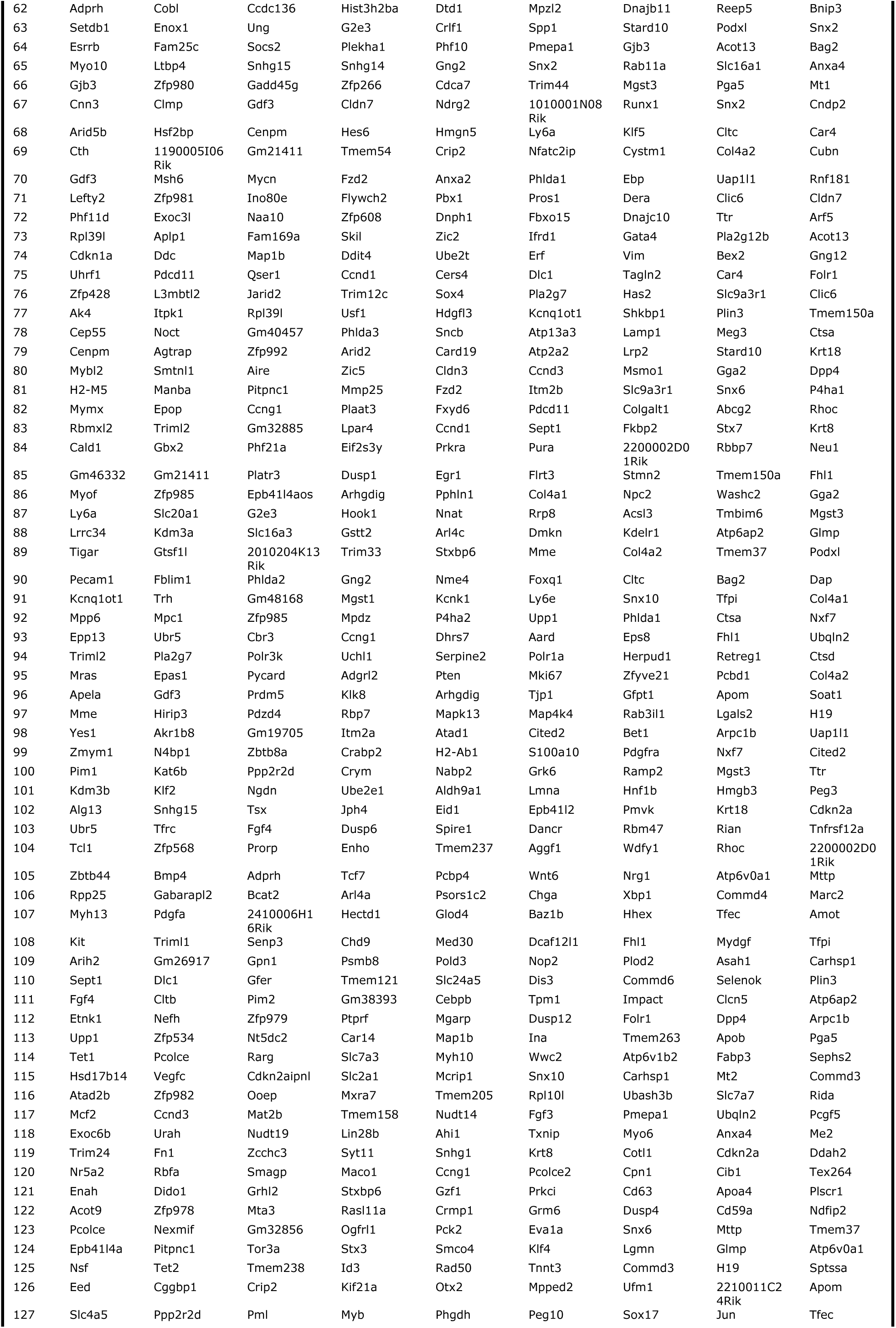
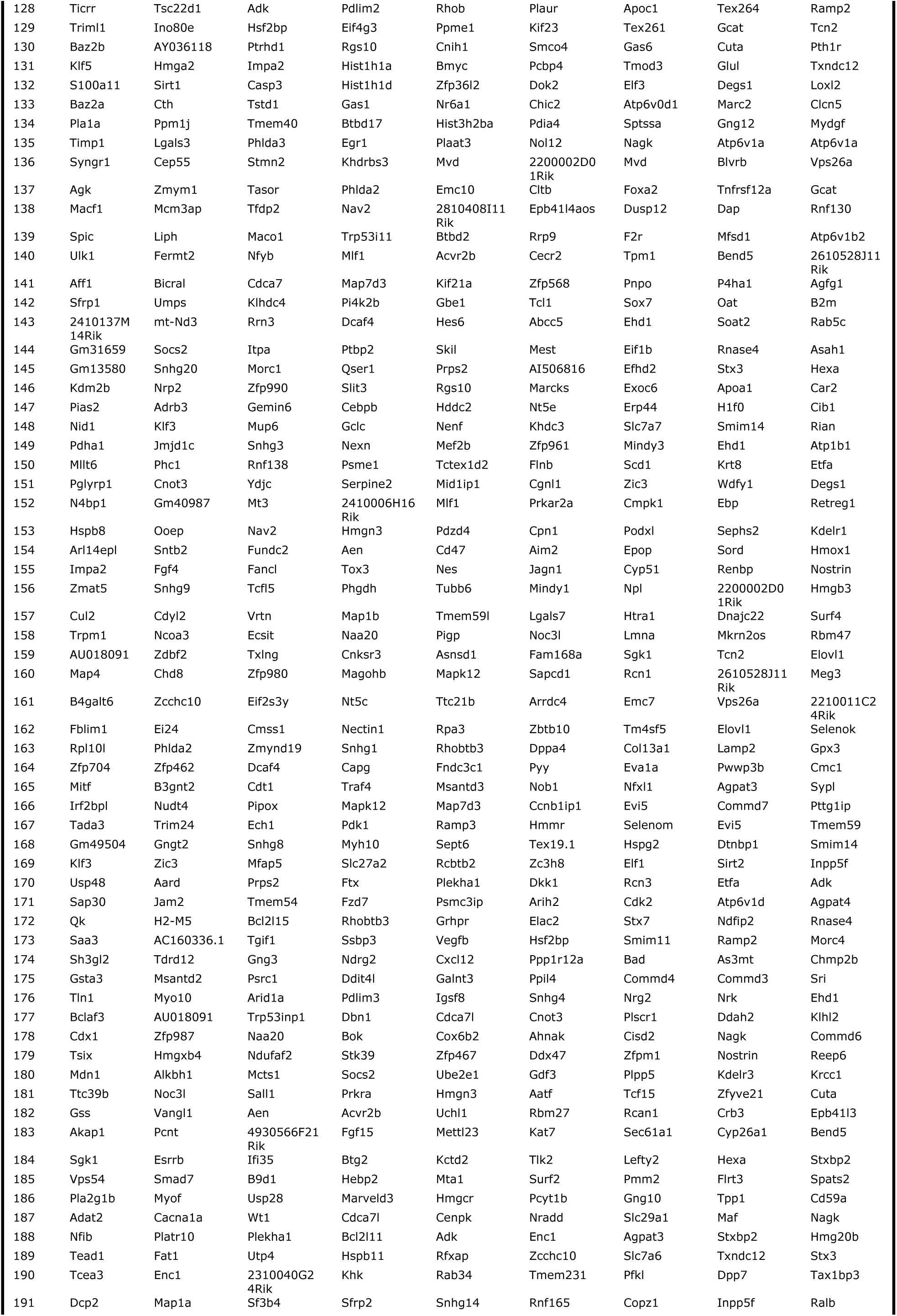
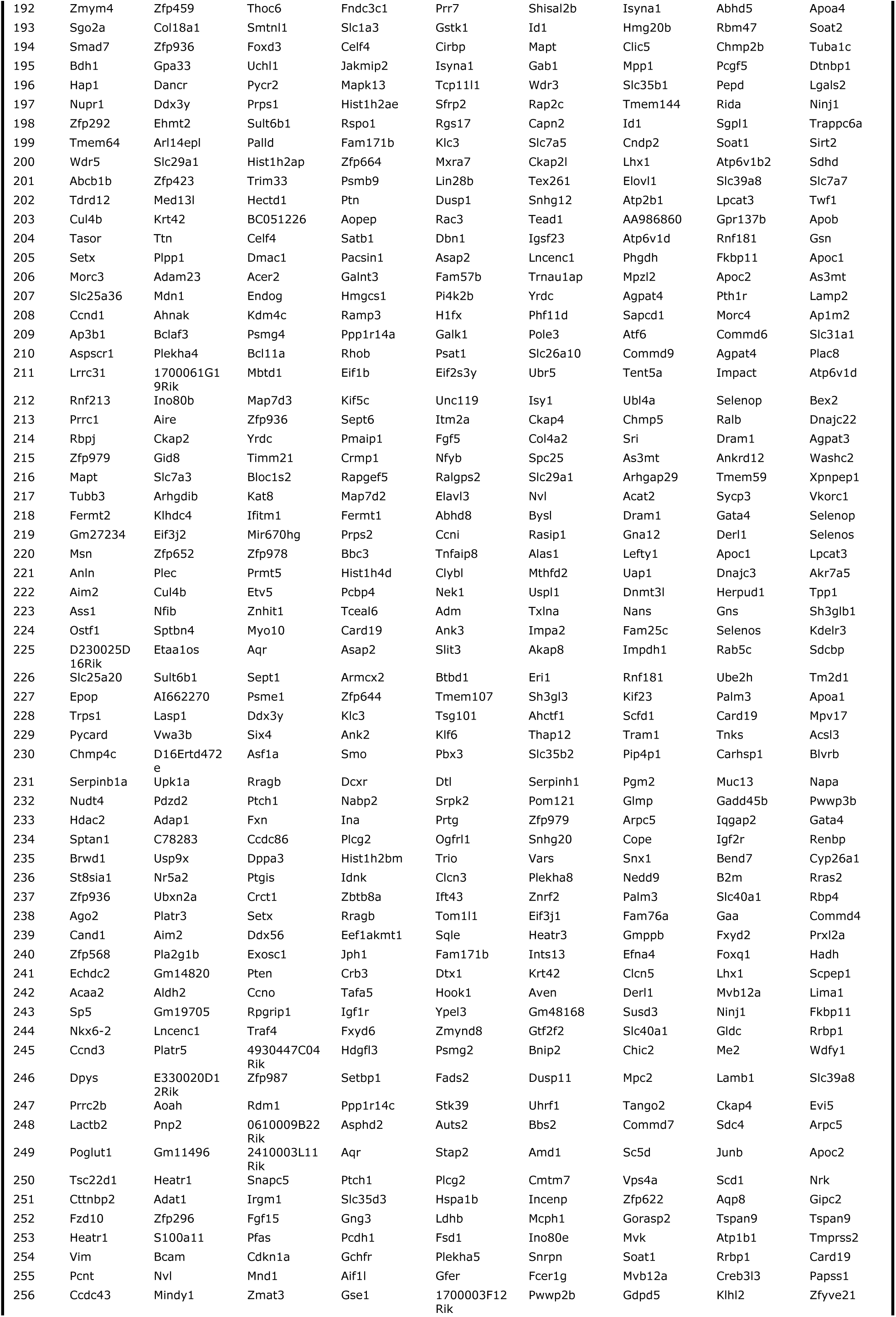
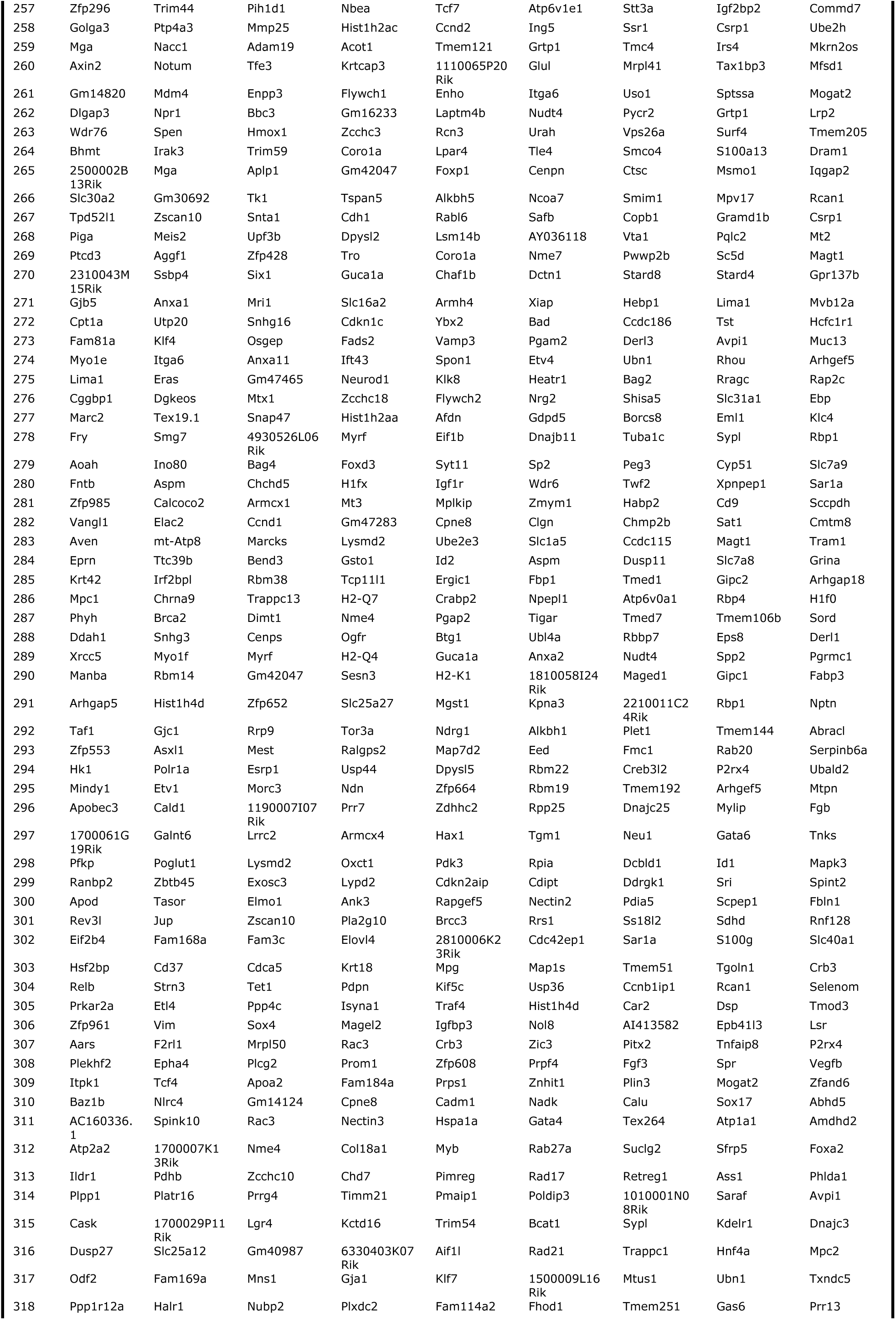
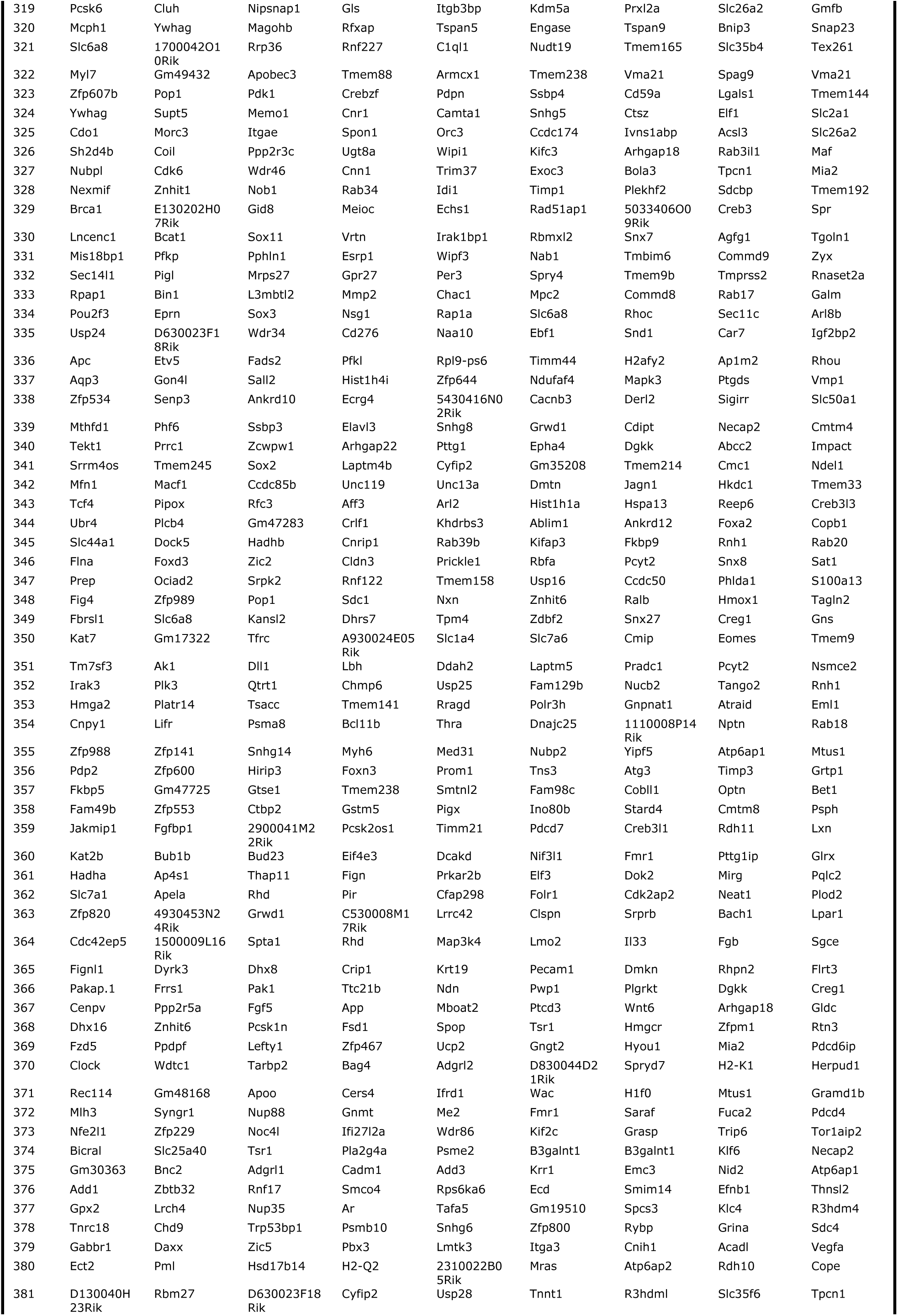
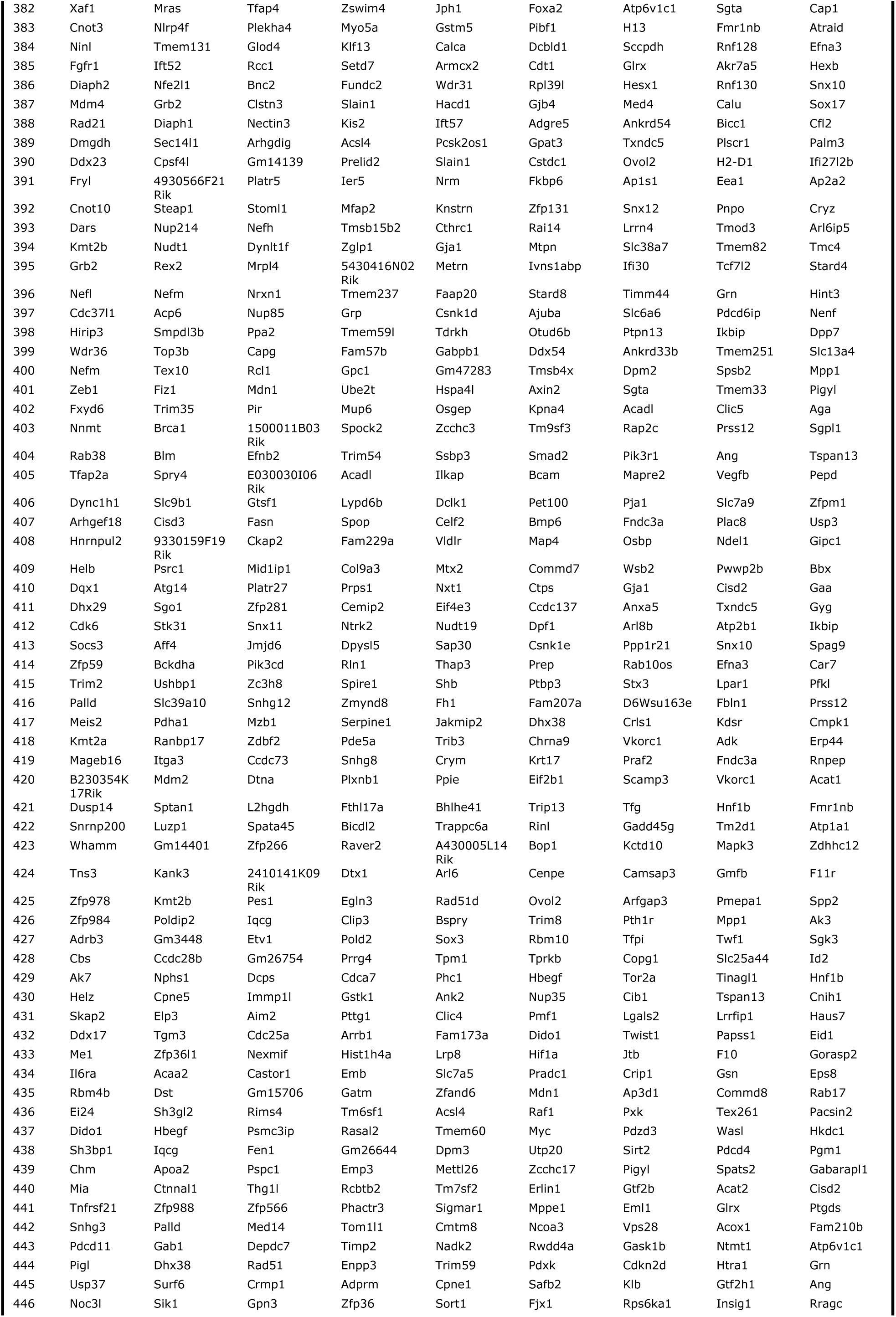
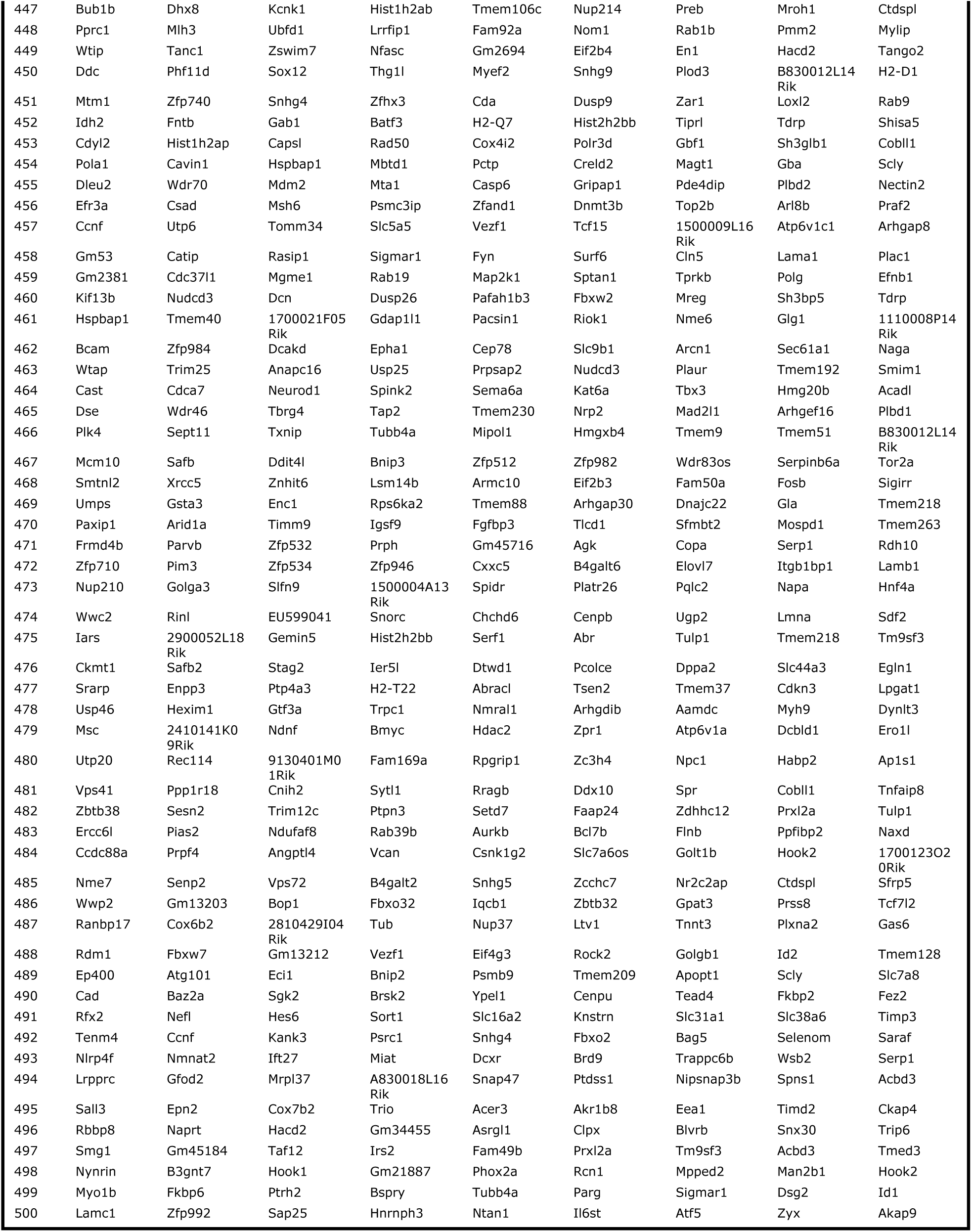
DEGs between different days of both lineages. For each group of cells, the top 500 DEGs are listed, which were also used for the comparison between ICM embryoid and embryo scRNA-seq data.

## Notes

### Competing Interest Statement

The authors have declared no competing interest.

